# Anti-tumor immunity relies on targeting tissue homeostasis through monocyte-driven responses rather than direct tumor cytotoxicity

**DOI:** 10.1101/2024.06.12.598563

**Authors:** Nicholas Koelsch, Faridoddin Mirshahi, Hussein F. Aqbi, Mulugeta Seneshaw, Michael O. Idowu, Amy L. Olex, Arun J. Sanyal, Masoud H. Manjili

**Affiliations:** Department of Microbiology & Immunology, Virginia Commonwealth University School of Medicine, Richmond, VA 23298, USA; Department of Internal Medicine, VCU School of Medicine, Richmond, VA 23298, USA; Stravitz-Sanyal Institute for Liver Disease and Metabolic Health, Richmond, VA 23298; College of Science, Mustansiriyah University, Baghdad, P.O. Box 14022, Iraq; Department of Pathology, VCU School of Medicine, Richmond, VA 23298, USA; VCU Massey Comprehensive Cancer Center, Richmond, VA 23298, USA; C. Kenneth and Dianne Wright Center for Clinical and Translational Research, Virginia Commonwealth University School of Medicine; VCU Institute of Molecular Medicine, Richmond VA 23298

**Keywords:** systems immunology, hepatocellular carcinoma, nonalcoholic fatty liver disease, cancer dormancy, inflammation, network medicine, stromal cells, Systems immunology, hepatocellular carcinoma, cancer dormancy, metabolic dysfunction–associated fatty liver disease

## Abstract

**Background:** Metabolic dysfunction–associated fatty liver disease (MAFLD) can progress to hepatocellular carcinoma (HCC), yet the immune mechanisms driving this transition remain unclear.

**Methods:** In a chronic Western diet (WD) mouse model, we performed single-nuclei RNA sequencing to track MAFLD progression into HCC and subsequent tumor inhibition upon dietary correction.

**Results:** Carcinogenesis begins during MAFLD, with tumor cells entering dormancy when HCC is mitigated. Rather than purely tolerogenic, the liver actively engages immune responses targeting myofibroblasts, fibroblasts and hepatocytes to maintain tissue homeostasis. Cytotoxic cells contribute to turnover of liver cells but do not primarily target the tumor. NKT cells predominate under chronic WD, while monocytes join them in HCC progression on a WD. Upon dietary correction, monocyte-driven immunity confers protection against HCC through targeting tissue homeostatic pathways and antioxidant mechanisms. Crucially, liver tissue response—not merely immune activation—dictates whether tumors grow or regress, emphasizing the importance of restoring liver tissue integrity. Also, protection against HCC is linked to a distinct immunological pattern, differing from healthy controls, underscoring the need for immune reprogramming.

**Conclusion:** These findings reveal the dual roles of similar pathways, where immune patterns targeting different cells shape distinct outcomes. Restoring tissue homeostasis and regeneration creates a tumor-hostile microenvironment, whereas tumor-directed approaches fail to remodel the TME. This underscores the need for tissue remodeling strategies in cancer prevention and treatment.

**Lay summary:** Our study challenges the traditional view that the liver is purely tolerant to immune responses, revealing that it actively regulates immunity to maintain tissue health. We found that liver cancer (HCC) begins during fatty liver disease (MAFLD) but can be halted if immune cells—especially monocytes—restore tissue integrity. Instead of focusing solely on killing tumors, effective immunotherapy should harness the body’s natural ability to repair the liver, creating an environment where cancer cannot thrive. This discovery paves the way for innovative treatments that promote immune-driven tissue regeneration as a strategy for cancer prevention and therapy.

## Introduction

Traditional interpretations of existing data suggest that immune responses are induced only in response to damage or infection, with the immune system being naturally tolerogenic to avoid reacting against harmless substances such as gut-derived nutrients in the liver ^1^. However, accumulating evidence challenges this view by demonstrating the presence of inflammatory immune responses under healthy conditions in the liver. The healthy liver maintains an active cytokine milieu, including pro-inflammatory cytokines such as IL-2, IL-7, IL-12, IL-15, TNF-α, and IFN-γ, as well as anti-inflammatory cytokines like IL-10, IL-13, and TGF-β, produced by the hepatic immune system ^2^. Additionally, activated NKT cells support hepatocyte proliferation and liver regeneration ^3^. Hepatic B and T cells also contribute to liver regeneration by producing lymphotoxin β (LTβ) ^4^, as the blocking of LTβR impedes liver regeneration after partial hepatectomy ^5^. Human hepatic CD141^+^ dendritic cells (DCs) are potent cytokine producers and activators of T cells ^6^. This homeostatic inflammatory immune response is essential for hepatic cell regeneration and tissue remodeling. However, the mechanisms by which hepatic immune responses are induced under healthy conditions or metabolic dysfunction–associated fatty liver disease (MAFLD) and hepatocellular carcinoma (HCC) remain elusive. The present study addresses this gap by the assessment of the hepatic immune system under healthy condition on a regular chow diet (CD) as well as during the progression or inhibition of HCC.

The immune system in each organ, including the liver, operates through a complex network of reciprocal cis and trans cellular interactions via ligand-receptor communications. Understanding this system requires a holistic perspective rather than focusing on individual cell types alone. This systems perspective is crucial for both the immunobiology of diseases and the development of novel immunotherapies. While reductionist approaches in immunology have advanced our knowledge of immune cell types, they fall short in uncovering the mechanisms of emergent collective functions of the organ-specific immune system, and in offering curative immunotherapies beyond prolonging the survival of cancer patients. Recent reviews have highlighted contradictory observations regarding the dual functions of innate and adaptive immune responses in tumor immune surveillance, inflammation-associated liver protection or liver damage, and promotion or inhibition of HCC ^7,8^.

Recent advancements in big data and computational algorithms have enabled the detection of cellular interactions as distinct networks, providing deeper insights into disease mechanisms. By adopting a systems immunology approach for understanding the progression of MAFLD to HCC, it has been demonstrated that dominant-subdominant relationships among hepatic immune cells shape immunological patterns from which collective functions emerge, distinct from the individual roles of each immune cell type ^9,10^. These dominant-subdominant interactions are analogous to "cell competition" or "cellular fitness," where winner cells survive and eliminate loser cells ^11,12^. In the present study, we focused on the molecular pathways of ligand-receptor interactions to uncover immunological patterns associated with dominant cell types and pathways involved in the progression or inhibition of HCC in the DIAMOND mouse model. This approach allows for a more comprehensive understanding of the mechanisms of MAFLD and HCC, as well as the development of more effective therapeutic strategies.

## Materials and Methods

### Animal model and experimental design

Diet-induced animal model of nonalcoholic fatty liver disease (DIAMOND) ^13^ were used in this study, which are an isogenic cross between C57BL/6J and 129S1/SvImJ mice. DIAMOND mice recapitulate many of the components seen in human MAFLD and HCC progression under conditions of Western Diet (WD Harlan TD.88137), a high fat diet (42% kcal from fat and 0.1% cholesterol) and ad lib administration of glucose (18.9 g/l) and fructose (23.1 g/l) in the drinking water ^13,14^, compared to no cholesterol and 5.8% kcal from fat in a standard chow diet (CD; Harlan TD.7012). This model is now being used for preclinical assessment of drugs before engaging in clinical trials. These studies have been reviewed and approved by the Institutional Animal Care and Use Committee (IACUC) at Virginia Commonwealth University on animal protocol number AD10001306. All methods were performed in accordance with the relevant guidelines and regulations. The use of this animal model allows the conduction of translational studies for the development of prognostic markers as well as treatment strategies for the prevention of human HCC. In brief, male mice were put on a standard CD for 40 weeks (n=2), WD, or underwent diet reversal (RD) after being on a WD. Only male mice were used in this study due to higher incidence of males developing HCC than female mice on a WD ^10^. Mice were on a WD for 40 weeks prior to the development of HCC and during MAFLD (WD.mf; n=2) or after development of HCC by 48 weeks (WD.t; n=3). Separate groups underwent diet reversal for additional 12 weeks, after 36 weeks of being on a WD, in which some mice developed tumors (RD.t; n=2) and some protected from the development of HCC and exhibited no tumors in the livers (RD.n; n=3) (Figure S1A).

### Hematoxylin and eosin staining

Formalin fixed paraffin embedded liver (FFPE) tissues were subjected to hematoxylin and eosin (H & E) stain using Tissue Tek Prisma Autostainer as previously described by our group ^15^. Histology slides were scanned at 40x magnification.

### Flow cytometry

Multicolor staining and flow cytometry analysis of T cells were performed as previously described by our group ^10^. Briefly, Fc blocker anti-CD16/32 Ab was used for all staining panels before using the T cell staining panel (CD8, CD4, CD44, CD62L). All reagents were purchased from Biolegend (San Diego, CA). All reagents were used at the manufacturer’s recommended concentration. Multicolor data acquisition was performed using a LSRFortessa X-20 (BD Biosciences) and a ImageStreamX Mark II Imaging Flow Cytometer (Millipore Sigma, Billaerica, MA). Data were analyzed using FCS Express v5.0 (De Novo Software; Glendale, CA). The FVS negative viable cells were gated on CD4+ or CD8+ T cells, and analyzed for CD44^+^CD62L^-//low^ T effector (Te), CD44^+^CD62L^+/high^ (Tcm) and CD44^-^CD62L^+^ T naïve (Tn) subsets.

### Quality control and filtering

Single nuclei RNA-seq data from all samples was provided by the Novogene company, in which nuclei were isolated from frozen mouse liver and tumor samples, and using 3’ single cell gene expression libraries after using three quality control methods for the library: Qubit 2.0 to test preliminary concentration, Agilent 2100 to test insert size, and Q-PCR to quantify library effective concentration precisely after Illumina Sequencing. Novogene performed 10x sequencing read processing and quality control by aligning N nucleotides of Read1s or Read2s against reference genome mm10-5.0.0 with STAR to filter and correct unique molecular identifiers (UMIs) so only confidently mapped, non-PCR duplicates with valid barcodes and UMIs were used for generation of gene-barcode matrixes for further analysis. Additionally, Novogene used Cell Ranger version 7.0.0 STAR aligner to perform splicing-aware alignment of reads to the genome, followed by transcript annotation GTF to bucket reads into exonic, intronic, and intergenic to determine if reads confidently align to the genome; a read is exonic if at least 50% of it intersects an exon, intronic if it is non-exonic and intersects an intron, and intergenic otherwise. Seurat version 4.4.0 ^16^ was used to handle all data and perform quality control and filtering metrics. In brief, samples were loaded into the VCU HPRC core clusters to mark data with group identifying labels (CD, WD.mf, WD.t, RD.t, RD.n), and filter cells with nFeature_RNA > 200 and < 5000, as well as for cells expressing < 5% mitochondrial associated genes. The threshold for nFeatureRNA > 200 and < 5000 is to ensure there are sufficient molecular transcripts in one cell and 5000 > may entail two cells in one run, whereas mitochondrial gene percentages higher than 5% may be indicative of dead/dying cells. The number of nuclei of cells sequenced per sample are as follows: CD1 = 7372, CD2 = 8132, WD.mf1 = 6160, WD.mf2 = 8469, MWD.t1 = 6457, MWD.t2 = 7083, MWD.t3 = 8872, MRD.t1 = 20000, MRD.t2 = 10522, MRD.n1 = 10861, MRD.n2 = 6049, MRD.n3 = 10688. After filtering dead/dying cells, each sample contained the following number of nuclei per sequencing sample: CD1 = 7231, CD2 = 7773, WD.mf1 = 5409, WD.mf2 = 7847, MWD.t1 = 2205, MWD.t2 = 4621, MWD.t3 = 3200, MRD.t1 = 19659, MRD.t2 = 9155, MRD.n1 = 10180, MRD.n2 = 4324, MRD.n3 = 9278. High Performance Computing resources provided by the High Performance Research Computing (HPRC) core facility at Virginia Commonwealth University (https://hprc.vcu.edu) were used for conducting the research reported in this work.

### Cell type annotation and quantification

Markers for major liver cell types specific for mice from liver focused data were extracted from the CellMarker 2.0 database ^17^ and compiled into one comprehensive list to annotate cells. The scSorter R program version 0.0.2 ^18^ was used in conjunction with these marker genes to annotate liver immune and nonimmune cell types such as B cell, T cell, DC, NKT, NK, neutrophil, monocyte, macrophage, endothelial, LSEC, stromal, HSC, fibroblast, myofibroblast, cholangiocyte, hepatocyte, and cancer cells in each group of samples individually (CD, WD.mf, WD.t, RD.t, RD.n), as this is more representative of clinical applications to individual patients. Following annotation, the cells were clustered and visualized in Seurat with UMAP after removing any cells classified as “Unknown” for subsequent analyses, representing a limitation of the scSorter program as only cells classified as known cell types are included. The number of “Unknown” removed per sample group ranged from 135 cells (0.89%) in the CD group, to 1365 cells (5.73%) in the RD.n group. The number of nuclei per sequencing sample after filtering dead/dying cells and removing “Unknown” classified include: CD1 = 7137, CD2 = 7732, WD.mf1 = 5101, WD.mf2 = 7691, MWD.t1 = 2196, MWD.t2 = 4445, MWD.t3 = 3133, MRD.t1 = 19553, MRD.t2 = 8474, MRD.n1 = 9581, MRD.n2 = 4287, MRD.n3 = 8549. Cellular annotations were confirmed by making heatmaps of marker genes specific for each cell type of interest (Figure S1B). Canonical correlation analysis (CCA) in Seurat version 4.4.0 was used on cells annotated by scSorter with 2000 variable features in the FindVariableFeatures function, followed by finding integration anchors, log-scaling the data, running PCA with 30 principal components, and running the UMAP function and finding neighbors and clusters with recommended default parameters. Further, quantification of the exact number of cells was performed in excel and normalized to 100% to show the composition of cell types in each group in the study, such as immune cells and nonimmune cells. Violin plots of specific marker genes in various cell type populations across groups were visualized and quantified in Seurat with the VlnPlot function. Assessments of specific genes such as Mki67 (Ki67), Tnfsf10 (TRAIL)/Tnfrsf10b (TRAILR2), FasL/Fas, Gzma, Gzmb, Prf1, Cxcl13, Ccl4, and IL-7 were performed in Seurat with the DotPlot function, and average transcript expression was quantified with AverageExpression function after filtering cell populations for only cells expressing the gene of interest (>0).

### Intercellular communication networks

CellChat version 2.1.1 (https://doi.org/10.1101/2023.11.05.565674) was utilized with default parameters (trimean approach requiring 25% of the cells in a population to express a specific ligand or receptor to be considered for statistical testing) to evaluate ligand and receptor (L-R) interactions amongst all annotated cell types. First, we performed this on all groups, and subsequently performed the same analysis with a “truncated mean” of 0.05 with the computeCommunProb function to evaluate lowly expressed immunologically relevant interactions in 5% of cells within each annotated cell type, as well as with a “truncated mean” of 0.75 to assess major communication networks in 75% of cells, however the capacity of filtering can only evaluate those in 50% of cells through the use of “truncated mean” of 0.50, as results from 0.75 were identical to 0.5. Additionally, by filtering L-R detection through the identifyOverExpressedGene function and filtering thresh.pc = 0.80 allows analysis of only L-R pairs expressed in 80% of each cell population. Higher threshold filtering such as 50% and 80% would be more biologically relevant, as the majority of the cell populations are expressing these molecules, while most cell-cell communication programs typically utilize a 10% population expression threshold for detection. Comparative CellChat analyses were also employed to address any changes in each cell type across groups, in which the sum of probability scores for cell types of interest were also quantified in excel to highlight the incoming and outgoing interaction strength in experimental groups. Analysis of differential number of interactions was performed to visualize the number of L-R interactions sent from one cell type to all others detected in one condition compared to the first, seen by red arrows indicating increased signaling events and blue arrows indicating decreased signaling events. Signaling pathway changes were analyzed similarly to compare major pathways utilized by a specific cell type in one condition compared to the first. Versions of additional dependencies for CellChat data visualization include NMF version 0.27 (https://github.com/renozao/NMF), circlize version 0.4.16 (https://github.com/jokergoo/circlize), and ComplexHeatmap version 2.20.0 (https://github.com/jokergoo/ComplexHeatmap).

### A <u>LI</u>gand-receptor <u>AN</u>alysis Fr<u>A</u>mwork (LIANA) utilization

LIANA ^19,20^ was utilized to confirm key interactions detected in the HCC tumor-bearing mice (WD.t), as well as mice that underwent diet reversal and developed HCC (RD.t), and those that did not (RD.n). The LIANA Rank Aggregate method was applied to each dataset individually and represents a consensus by integrating the predictions of individual methods from multiple programs L-R detection methods (CellPhoneDB, Connectome, log2FC, NATMI, SingleCellSignalR, Geometric Mean, scSeqComm, and CellChat). The results were filtered to show L-R interactions detected in 50% CellChat analyses that were also present in the LIANA consensus database (Plg-Pard3, Pros1-Mertk, Vtn-Itgav_Itgb1, Vtn-Itgav_Itgb5, Angptl4-Sdc2, Angptl4-Sdc4, and Hgf-Met). Specificity rank shows how specific an interaction is to a pair of cell types, in which larger the dot size, the more specific to a pair of cell types it is. Magnitude rank shows the corresponding strength of that detected LR interaction, with yellow being stronger and blue being weaker. These consensus metrics are generated in LIANA across all methods using Robust Rank Aggregation ^21^, that can be interpreted as probabilistic distributions for interactions that are more highly ranked ^19,20^. This analysis was performed in Python 3.11.6 and High Performance Computing resources provided by the High Performance Research Computing (HPRC) core facility at Virginia Commonwealth University (https://hprc.vcu.edu) were used for conducting the research reported in this work. In addition, the version of programs utilized in python include: LIANA 1.4.0, scanpy 1.9.6, anndata 0.11.2, pandas 2.2.3, and plotnine 0.14.5.

### Statistical analysis

CellChat identifies differentially expressed signaling genes via Wilcoxon rank sum test filtering for those at a 0.05 significance level, followed by averaging gene expression across a given cell group, generation of intercellular communication probability values for each L-R interaction, and finally identifying statistically significant communications through permutation tests and recalculating communication probability between cell groups (18). Chord diagrams of signaling directionality were further filtered to only include L-R signaling interactions that were at or below a p-value of 0.01. Quantification of gene expression levels, such as Hnf4α and Ki67, were averaged across replicates and calculated standard error mean by the standard deviation of expression levels in sample replicates divided by the square root of the number of sample replicates. A two-tailed Student’ *t*-test was used to analyze flow cytometry data, and a *p*-value of ≤ 0.05 was considered statistically significant. The analysis focuses on variability between cells rather than entire organisms, minimizing the impact of inter-animal variability and allowing meaningful conclusions from fewer animals per group. For discovery-driven projects exploring individuality, snRNA-seq inherently supports single-animal resolution by revealing cell-type-specific and condition-specific molecular signatures. Advanced computational tools and algorithms for snRNA-seq enable robust analysis of small datasets, reducing the need for large animal cohorts while maintaining statistical power. snRNA-seq provided detailed single-nucleus transcriptional profiles, capturing rare cell populations and subtle molecular variations that revealed individual differences during WD (Figure S2). This underscores the need for individual-based analysis to establish functional cohorts, enabling more precise insights from smaller, focused studies in the future

### Data and code availability

All code and gene lists for sorting cells are available on GitHub (https://github.com/koelschnj/Systems-Approaches-Understanding-Liver-Cell-Networks-MAFLD-HCC-DietReversal) and is publicly available on the following links through the Gene Expression Omnibus (GEO) ^22^: https://www.ncbi.nlm.nih.gov/geo/query/acc.cgi?acc=GSE225381 and https://www.ncbi.nlm.nih.gov/geo/query/acc.cgi?acc=GSE279124. The CD and WD.mf datasets analyzed during the current study are available in the GEO repository, GSE225381. The WD.t, RD.t, and RD.n datasets generated and analyzed during the current study are available in the GEO repository, GSE279124.

### Materials availability

DIAMOND mice used in this study will be provided, upon request to the lead contact, and may require fulfillment of an MTA. This study did not generate new unique reagents.

## Results

### Carcinogenic events take place during MAFLD and remain dormant during protection against HCC

The livers of male DIAMOND mice that were on a CD or WD or reversal of a WD to a CD for 12 weeks starting from 36 weeks of being on a WD were examined and subjected to hematoxylin-eosin (H&E) staining and single nuclei RNA sequencing (snRNAseq) analysis when animals reached 13 months of age (Figure S1A). There was no macroscopic tumor based on visual inspection, or microscopic tumor, based on histology, detectable in the liver of animals during MAFLD at 11 months of age (WD.mf) or protection against HCC following dietary correction at 13 months of age (RD.n) (Figure 1A). However, cell annotation and UMAP clustering of immune and nonimmune cells by using marker genes from the CellMarker2.0 database (Figure S1B) revealed a cancer fraction in the liver (Figure 1B), characterized by the oncogenic glypican-3 (GPC3) and alpha-fetoprotein (AFP), as well as the tumor suppressor retinoblastoma 1 (Rb1) and hepatocyte nuclear factor 4 alpha (Hnf4α) ^23^, separating them from hepatocytes (Figure 1C). In fact, the hepatocyte fraction mainly expressed the tumor suppressor genes Hnf4 or Rb1, whereas cancer fraction expressed the oncogene markers GPC3 and AFP as well as the tumor suppressor genes (Figure 1C). Further filtration based on the expression of GPC3 oncogene transcript, which is more sensitive than AFP in the diagnosis of liver cancer ^24^, revealed high expression of all tumor markers in cancer fraction of the WD.t group followed by three markers in the WD.mf and RD.t groups whereas the RD.n group showed lower expression of the oncogenes and higher expression the Hnf4α tumor suppressor gene (Figure 1D), The CD group showed only two markers, GPC3 and Rb1 (Figure 1D). Among these groups, the highest level of Hnf4α transcript was detected the CD group followed by the RD.n group, whereas the lowest levels of Hnf4α transcript were detected in the WD.mf and RD.t groups (Figure 1E). Although the presence of cancer fraction in the CD group was minimal (Figure 1B), this may be due to age-associated carcinogenesis events which could be controlled by highest level of the Hbf4α tumor suppressor gene. Further analysis of the CD, WD.mf and RD.n groups showed differential average transcript for Ki67 expression between the cancer and hepatocyte fractions. While hepatocytes showed high level of Ki67 transcripts (>5) in the WD.mf group compared to intermediate levels (between 2-5) in the CD and RD.n groups, cancer fractions showed Ki67 negative quiescent cells in the CD group and Ki67^low^ (<2) indolent cells in the WD.mf and RD.n groups (Figure 1F). We have already reported that indolent dormant cells establish cancer as opposed to quiescent cells remaining dormant ^25,26^. It is yet to be determined whether such malignant events signify tumor dormancy or the presence of cancer stem cells. We have previously detected such tumor dormancy in the FVBN202 transgenic mouse model of breast cancer ^25^.

**Figure 1.**
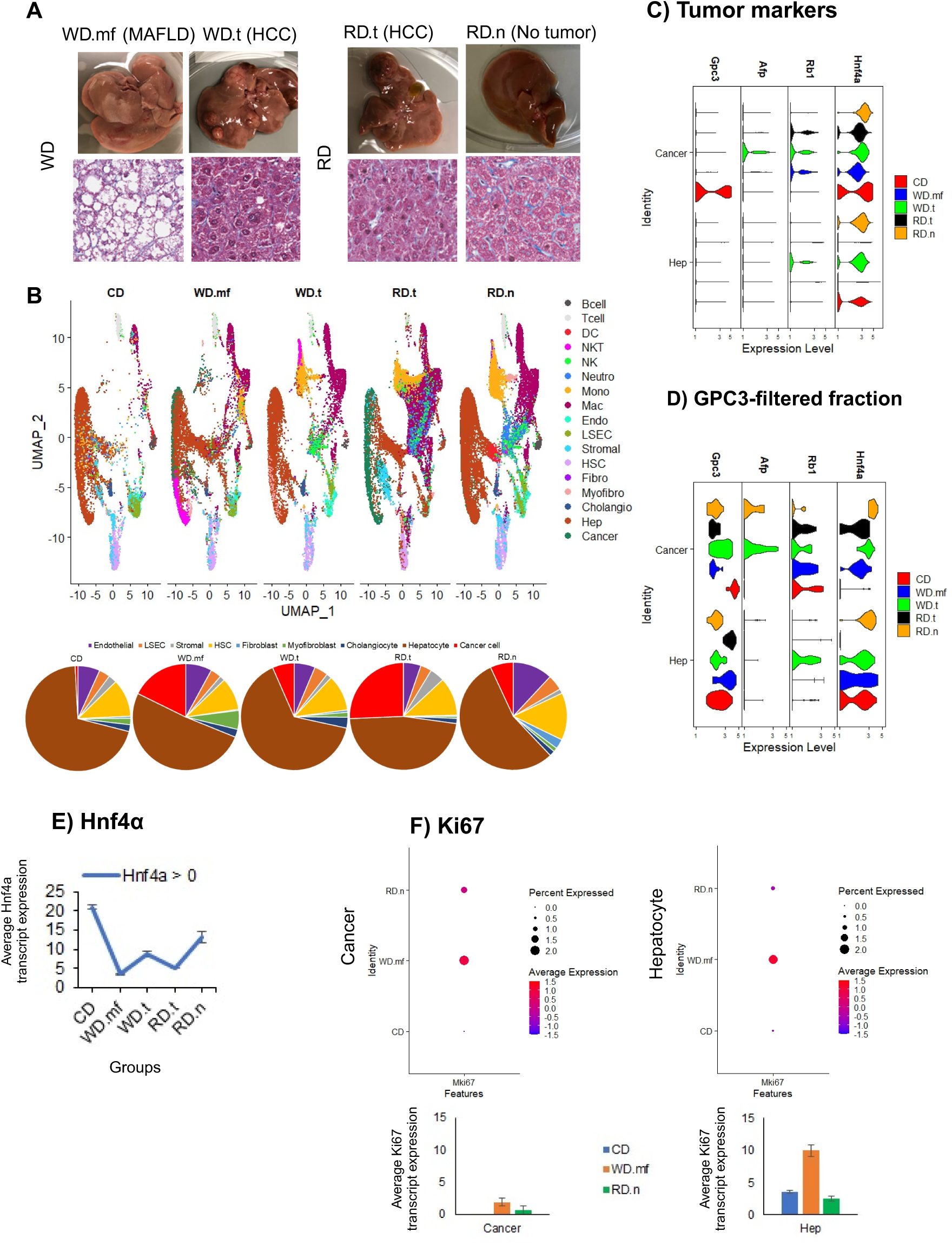
Carcinogenic events take place prior to HCC and remain dormant during recovery from HCC. A) Representative liver images and histological Hematoxylin and Eosin (H&E) staining from mice with MAFLD (WD.mf; n=2), WD-induced HCC tumors (WD.t; n=3), and those that underwent dietary intervention and showed HCC tumors (RD.t; n=2) or those that did not (RD.n; n=3). B) UMAP of hepatic immune and nonimmune cells (upper) annotated by scSorter after removing all cells classified as “Unknown”, and quantified nonimmune cell proportions (lower); legend abbreviations include dendritic cell (DC), Neutrophil (Neutro), Monocyte (Mono), Macrophage (Mac), Endothelial cell (Endo), Liver sinusoid endothelial cell (LSEC), hepatic stellate cell (HSC), Fibroblast (Fibro), Myofibroblast (Myofibro), Cholangiocyte (Cholangio), and Hepatocyte (Hep). C) Violin Plots showing log-transformed average transcript expression of marker genes in cancer cell and hepatocyte populations such as Hnf4α, Rb1, Afp, and Gpc3. D) Violin Plot showing log-transformed average transcript expression in the Gpc3-filtered fraction of hepatocyte and cancer cells for each group. E) Quantification of the average transcript expression level of Hnf4α+ cells in the cancer cell population in each group with standard error mean calculated and applied as error bars across replicates. F) DotPlots showing the average expression level and percentage of cell populations expressing Mki67 (Ki67) (upper) in cancer cell (left) and hepatocyte (right) populations for the CD, WD.mf and RD.n groups, along with the average transcript expression in both cell populations after filtering for cells expressing Ki67 > 0 (lower) with standard error mean calculated and applied as error bars across replicates.

### Hepatic structural cells induce immune responses that participate in liver homeostasis

In order to determine whether liver is a tolerogenic organ, the crosstalk between hepatic structural cells and immune cells during liver tissue homeostasis under normal physiological conditions was investigated. We analyzed the livers of one-year-old mice that had been on a regular CD for 40 weeks, using scSorter and CellChat Analyses. A quantitatively predominant adaptive immune cells, T and B cells, was detected (Figure 2A). However, the receptor-ligand interaction analyses revealed a lack of immune cells engagement at the 80% threshold along with predominant hepatocytes sending homeostatic PARs (Protease-activated receptors) signal to structural cells (Figure 2B). Notably, 25-50% of macrophages were activated by myofibroblasts and cholangiocytes via Gas6-Mertk/Axl signaling, promoting efferocytosis (Figure 2C-D) ^27,28^. Additionally, 50% of macrophages and monocytes contributed to liver homeostasis through the IGF-IGFR1 (Insulin Growth Factor Receptor 1) and FN1 (Fibronectin)-Sdc4 (Syndecan 4) pathways, respectively (Figure 2C). IGF-1 is involved in normal glucose homeostasis in the liver ^29^, the activation of myofibroblasts and HSCs, as well as modulation of DC maturation ^30^. The Sdc4 signaling is involved in liver tissue integrity and cell-matrix remodeling ^31^. Also, 25% of hepatic cells were involved in the activation and modulation of immune responses through immune Gal9 (Galectin 9)-CD44/CD45, Spp1 (Secreted Phosphoprotein 1)-CD44/VLA4 (Very Late Antigen 4 or Itga4+Itgb1), Laminin-CD44/Dag1, Collagen-CD44, FN1-CD44/VLA4, Cholesterol-Rora (Retinoic acid-related orphan receptor alpha) and anti-inflammatory DHEAS (dehydroepiandrosterone sulfate)-Pparα/δ (Peroxisome proliferator-activated receptor alpha or delta) that targeted PPARα ^32,33^, while 25% of immune cells participated in liver homeostasis through laminin and VTN (Vitronectin) (Figure 2D). Galectin 9 has been reported to induce T cell activation through the engagement with the dominant pathway CD45 that regulates signaling thresholds by dephosphorylating components of the Src kinase family, and LcK-dependent calcium mobilization in peripheral CD4^+^ T cells ^34^. Also, the Gal9-CD44 interaction enhances stability and function of adaptive Tregs ^35^. Finally, Galectin 9 binds IgM-BCR to regulate B cell signaling ^36^. Targeting Dag1 in the laminin pathway can activate phospholipase C gamma (PLC-γ) downstream of the TcR-CD1d in NKT cells ^37^.

**Figure 2.**
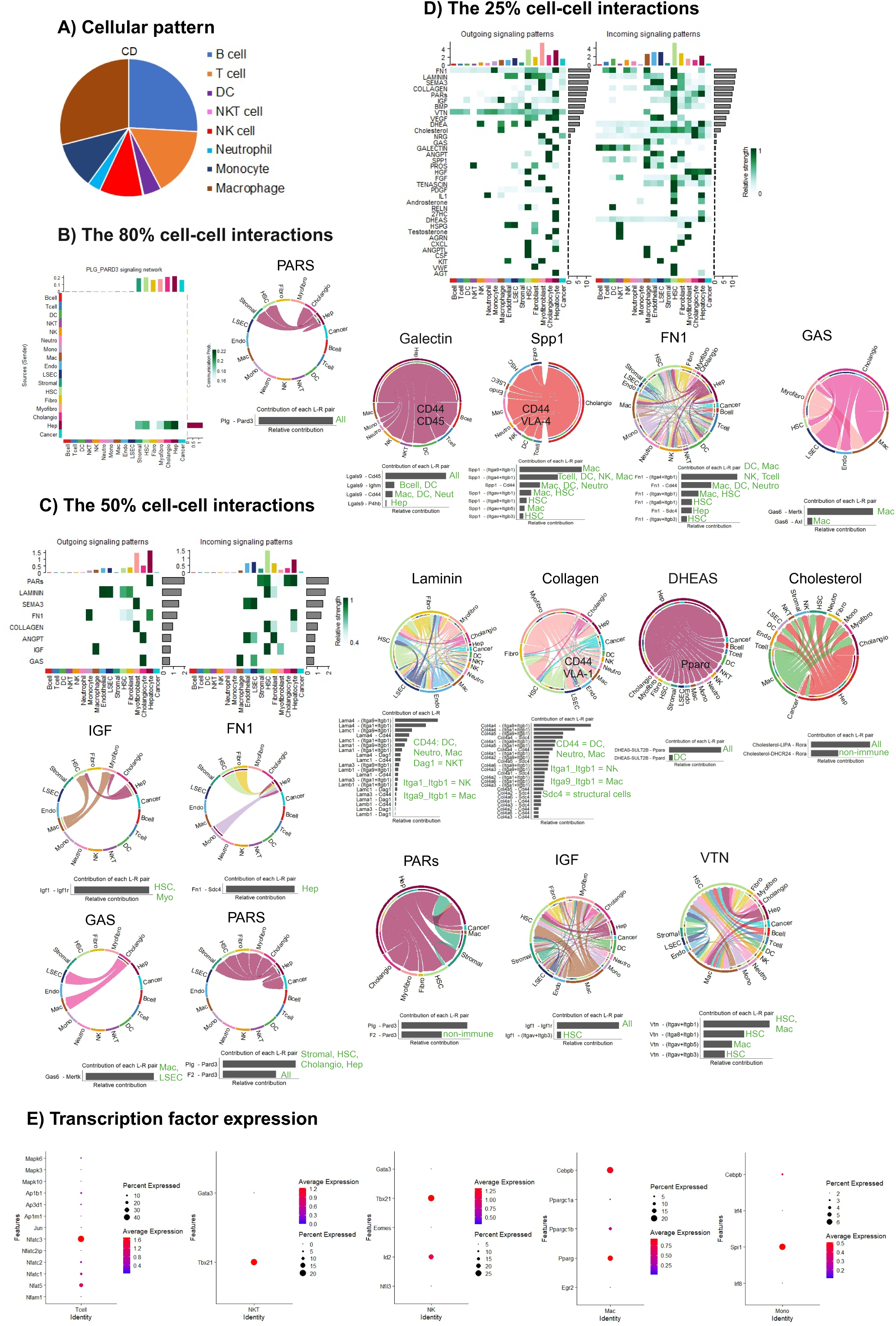
Hepatic cells induce macrophage and monocyte-dominated immune responses through integrins for liver tissue integrity and homeostasis. A) Immune cell proportions quantified and normalized to 100% in the CD group (n=2). B) CellChat analysis heatmap showing results from 80% threshold analysis for Ligand (L) and Receptor (R) interactions in the CD group and representative chord diagram showing signaling directionality in the PARs pathway and the L-R contributions. C) CellChat analysis heatmap portraying results from 50% threshold analysis for L-R interactions in the CD group (upper), and representative chord diagrams showing signaling directionality in the IGF, FN1, GAS, and PARs dominant pathways and their L-R contributions (lower). D) CellChat analysis heatmap portraying results from 25% threshold analysis of L-R interactions in the CD group (upper), and the representative subdominant signaling chord diagrams detected for Galectin, Spp1, FN1, GAS, Laminin, Collagen, DHEAS, Cholesterol, PARs, IGF, and VTN pathways and their L-R contributions (lower). E) DotPlots showing the percent expression in cell populations and log-normalized average transcript expression of transcription factors involved in activation of T cells (left), NKT cells (middle left), NK cells (middle), Macrophages (middle right), and Monocytes (right). Green text in L-R contributions denotes the cell type expressing receptors and Itga4+Itgb1 denotes VLA-4.

Analysis of the nuclear transcription factors associated with immune cell activation showed presence of activated T cells (NFAT), NKT and NK cells (T-bet and Id2), macrophages (pro-inflammatory C/EBPβ and anti-inflammatory PPARγ), and monocytes (SPI1/PU.1) (Figure 2E). Along this line, flow cytometry analysis of the hepatic T cells showed presence of CD44^+^/CD62L^-/low^ CD4^+^ Te and CD8^+^ Te cells in the liver (Figure S3A). Consistent with our observations, it was reported that both CD4^+^ and CD8^+^ T cells, but not γδ T cells, are required for normal liver regeneration through lymphotoxin production such that RAG1^−/−^ mice show extensive hepatic injury following partial hepatectomy ^38^. These data suggest the tissue-based direct activation of the immune cells as well as participation of activated immune cells in liver homeostasis during normal condition.

Regarding immune cell cytotoxicity, around 10% of NK cells and 20% of NKT cells were involved in TRAIL-mediated turnover of 6% of fibroblasts (Figure S3B, upper panels). Additionally, 15% of NK cells and 5% of T cells were involved in Fas-L-mediated turnover of 20% of fibroblasts, endothelial cells, and LSEC (Figure S3B, lower panels). Other cytotoxic pathways included 10-15% of NK cells expressing granzyme A or B, and 20% of NKT cells expressing perforin (Figure S3C). All these hepatic structural cells were found to express MHC class I molecules, primarily H2-Q10 and H2-K1 (Figure S3D).

### Qualitative cell-cell interaction networks rather than quantitative cell type dominance determine the progression or inhibition of HCC

The hepatic cells were analyzed for the identification of cell types of which 50% participated in ligand-receptor interactions network. The highest number and strength of ligand-receptor interactions were evident during WD as well as protection against HCC following diet reversal (Figure 3A). In order to identify cell types that dominate the communication networks under each condition, cell type ratios and cell-cell interactions were analyzed. In all groups, hepatocytes and macrophages were quantitively dominant cell populations compared to other nonimmune and immune cell types (Figure 3B, left panels). However, the ligand-receptor interactions showed predominant cancer cells and NKT cells in sending (outgoing) and receiving (incoming) signals during MAFLD, with monocytes joining them in predominant cell-cell communication network during HCC on a WD. Maintenance of monocytes dominance during diet reversal was associated with protection against HCC (Figure 3B, left panels). Although there are contradictory reports on the role of NKT cells as well as T cells, macrophages, and monocytes in promoting or inhibiting HCC ^39–46^, our data suggest that NKT cell signaling network became dominant in response to WD, as they disappeared following dietary correction. Also, monocytes were active in sending signals for protecting from HCC in the WD.t group, but they did not succeed because of chronic intake of a WD; however, their signaling retention during dietary correction protected animals against HCC, whereas disappearance of their signaling network resulted in HCC progression following dietary correction. These data suggest that elevated cell-cell communication network, but not the abundance of infiltrated cells, may be of host-protective nature such that failures in host protection were either due to the overload of chronic WD consumption or diminished levels of cell-cell interactions in the RD.t group.

**Figure 3.**
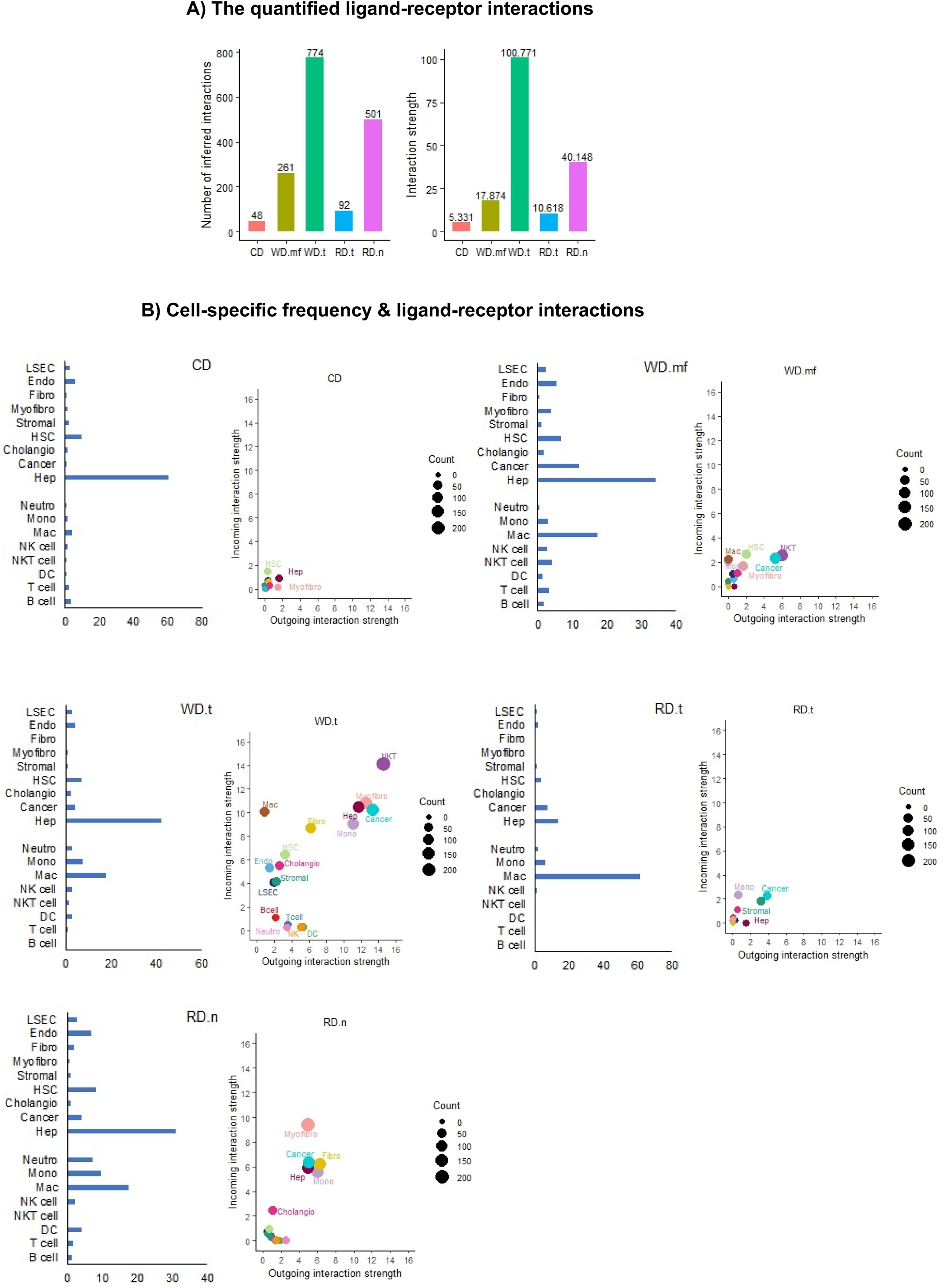
The number and strength of ligand-receptor interactions rather than cell type frequency determine the progression or inhibition of HCC. A) Comparative 50% threshold CellChat analysis quantifying the number of inferred interactions and overall interaction strength in each group. B) Cell-specific frequency (left panels) and comparative analysis in each group (right panels) showing overall incoming and outgoing interaction strength values for each cell type to evaluate major contributors to signaling networks. LSEC, liver sinusoidal endothelial cell; Endo, Endothelial cell; Fibro, fibroblasts; Myofibro, myofibroblasts; Stromal, stromal cell; HSC, hepatic stellate cell; Cholangio, cholangiocytes; Cancer, Cancer cell; Hep, hepatocytes; Neutro, neutrophils; Mono, monocytes; Mac, macrophage; DC, dendritic cell.

Among nonimmune cell-cell communication network, cancer cells were dominant during MAFLD or HCC following dietary correction, with myofibroblasts, hepatocytes and fibroblasts joining them during HCC on a WD or protection against HCC (Figure 3B, right panels). These data suggest that cancer cells, both as dormant and proliferating tumor, dominate the ligand-receptor interactions network regardless of their quantity.

### NKT cells orchestrate homeostatic immune responses in cancer-dominated networks during WD-induced MAFLD progression

Since cell-cell interaction networks were found to be more important than quantitative abundance of cell types, we sought to determine molecular pathways that dominate cell-cell interactions during the progression of MAFLD. Although pattern of cellular distribution showed higher proportion of hepatocytes and macrophages (Figure 4A), analysis of signaling pathways revealed NKT cells and cancer cells dominating 80% of the ligand-receptor cellular interactions (Figure 4B). They were involved in the activation of all immune cells, but NKT cells, through the LcK activating Gal9-CD45 pathway as well as the activation of monocytes through FN1 interacting with the co-stimulatory VLA-4 (Itga4+Itgb1) ^47^ (Figure 4B). They were also involved in the modulation of NKT cells and hepatic structural cells through the cholesterol-RoRα pathway which induces fatty acid oxidation and modulation of inflammation in immune cells ^48^ during immune responses ^49^ as well as affecting liver structural cells for hepatic lipid homeostasis ^50^. Such dominant NKT cell response was associated with the expression of CD1d on hepatic cells, particularly on cancer cells (Figure 4C). They also participated in PARs signaling that facilitates cell polarity and tight junctions and is essential for maintaining the integrity of the liver tissue, and facilitating hepatocytes repair and regeneration process during liver injury. Though 80% of hepatocytes lost this homeostatic pathway while cancer cells received PARs signaling during MAFLD (Figure 4B). Analysis of 50% of cells contributing to the ligand-receptor interaction network showed similar pattern with a broader receptor targeting of the structural cells and immune cells (Figure 4D). Cancer cells and NKT cells participated in the activation of all immune cells through Gal9-CD45/CD44 pathway, with cholangiocytes and HSCs joining them in activating macrophages and monocytes through FN1-CD44/VLA-4 pathway (Figure 4D). Additional homeostatic signaling pathways contributing to the activation of macrophages, monocytes and NKT cells included laminin and collagen (Figure S4). NKT cells joined cancer cells in targeting endothelial cells and LSEC to promote vascularization through VEGF (Figure S4). NKT cells also targeted myofibroblasts homeostasis by manifesting anti-fibrotic function via testosterone and pro-fibrotic function via FGF, PDGF and 27HC pathways, as well as targeting cancer fraction promoting HCC via EGF and ANGPTL pathways (Figure S4).

**Figure 4.**
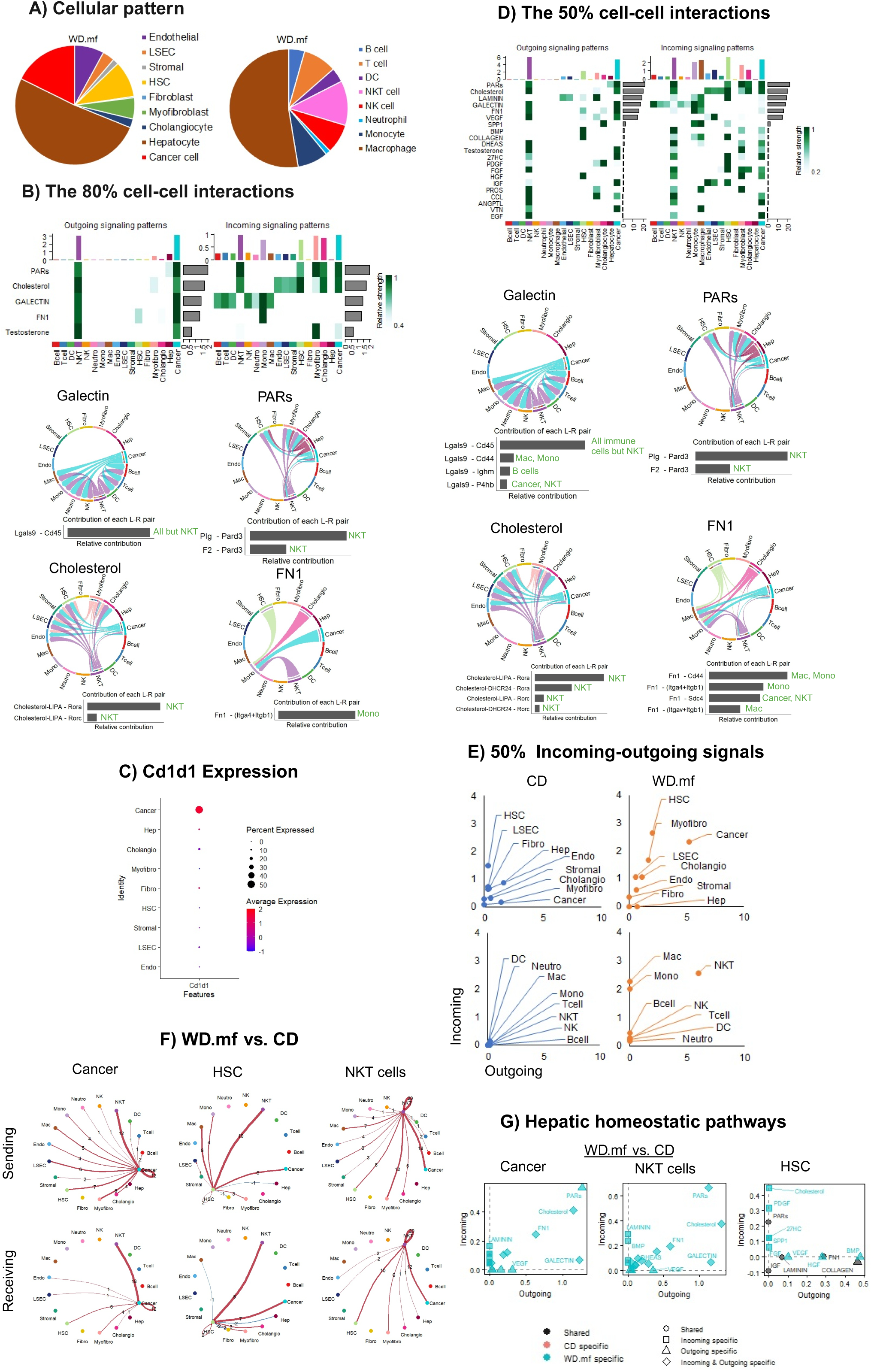
NKT cells orchestrate homeostatic immune responses over cytotoxicity in cancer-dominated networks during WD-induced MAFLD progression. A) Immune and nonimmune cell proportions quantified and normalized to 100% in the WD.mf group (n=2). B) CellChat analysis heatmap from 80% threshold results (upper) and representative chord diagrams showing signaling directionality in Cholesterol, PARs, Galectin, and FN1 pathways and their L-R contributions (lower). C) DotPlot showing the percentage of cell population expression and log-normalized average transcript expression of Cd1d1 in structural cells in the WD.mf group. D) CellChat heatmap analysis of 50% cell-cell interactions portraying all detected L-R signaling pathways (upper), and representative chord diagrams of signaling directionality in Cholesterol, PARs, Galectin, and FN1 pathways and their L-R contributions (lower). E) Quantification of cell-specific incoming and outgoing interaction strength in 50% CellChat analyses for nonimmune (upper panel) and immune cells (lower panel) in the CD and WD.mf groups. F) The differential number of signaling events sent (upper row) and received (lower row) detected in HSCs, cancer cells and NKT cells encompassing 50% of cell-cell interactions during MAFLD (WD.mf) compared to the CD group, where red arrows show increased number of signaling events in WD.mf and blue arrows show decreased events compared to CD, and thicker red or blue arrows show higher increases or lower decreases in signaling events across groups. G) Signaling pathway changes in 50% of cell-cell interactions in the WD.mf group compared to the CD group for HSCs, cancer and NKT cells. LSEC, liver sinusoidal endothelial cell; Endo, Endothelial cell; Fibro, fibroblasts; Myofibro, myofibroblasts; Stromal, stromal cell; HSC, hepatic stellate cell; Cholangio, cholangiocytes; Cancer, Cancer cell; Hep, hepatocytes; Neutro, neutrophils; Mono, monocytes; Mac, macrophage; DC, dendritic cell. Green text in L-R contributions denotes the cell type expressing receptors focused mainly on immune cells, and Itga4+Itgb1 denotes VLA-4.

Since signaling network dominance by hepatocytes during CD shifted to predominant NKT cell and cancer cell signaling during MAFLD, we performed a comparative analysis of signaling directionality between these groups. During MAFLD, the interaction networks accounting for 50% of cells were dominated by cancer, HSCs and NKT cells (Figure 4E). All these cells received highest number of signals from NKT cells and then cancer cells compared to the CD group (Figure 4F). In cancer cells and NKT cells, the significantly increased signals included reciprocal PARs, cholesterol, Gal9 and FN1 (Figure 4G). Fifty percent of all other immune cells did not change the number of outgoing signals during MAFLD compared to those in the CD group, but they received highest number of signals from NKT cells and cancer cells (Figure S5A). Among 50% of other structural cells, all but hepatocytes and fibroblasts were impacted mainly by NKT cells and cancer cells (Figure S5B). Despite NKT cells dominating the signaling network, they lacked cytotoxic function such as Granzyme A and B, perforin, Fas-L and TRAIL (Figure S6A-B). Ten percent of NK cells and T cells were involved in the turnover of 3-6% of cholangiocytes, myofibroblasts, and fibroblasts, while 10-15% of NK cells and T cells participated in Fas-mediated apoptosis of 30% of cancer cells and the turnover of 20-30% of myofibroblasts, LSECs, and endothelial cells (Figure S6B).

### Cancer cells, myofibroblasts and hepatocytes drive NKT- and monocyte-led immune responses via homeostatic pathways during HCC on a WD

While hepatocytes and macrophages remained quantitively dominant cell types during HCC on a WD (Figure 5A), 80% of the ligand-receptor interaction network was dominated by monocytes and NKT cells as well as cancer, myofibroblasts and hepatocytes, sending six homeostatic signals (Figure 5B). Among these signals, all immune cells followed predominant monocytes and NKT cells in sending PARs and FN1 signals (Figure 5B). In immune cells, PARs signaling promotes phagocytosis in macrophages and monocytes by facilitating the formation of the phagocytic cup, and also supports the formation of the immune synapse, enabling efficient cell-cell communication and T cell activation. The FN1-SDC4 pathway induces myofibroblasts and fibroblasts to promote tissue healing as well as proinflammatory cytokine secretion by NKT cells and monocytes. Monocytes and NKT cells were the only immune cells that sent DHEAS, 27HC, and FGF signals to themselves, hepatic structural cells, and cancer cells (Figure 5B). They also delivered VEGF to endothelial cells and LSEC (Figure 5B). To this end, DHEAS, 27HC, and FGF inhibit their inflammatory functions. These predominant signaling pathways engaged additional receptors, such as Itgav-Itgb1, CD44, PPARγ, and FGFR4, accounting for 50% of cell-cell interactions (Figure 5C). These data suggest simultaneous activation and inhibition of the immune responses during HCC.

**Figure 5.**
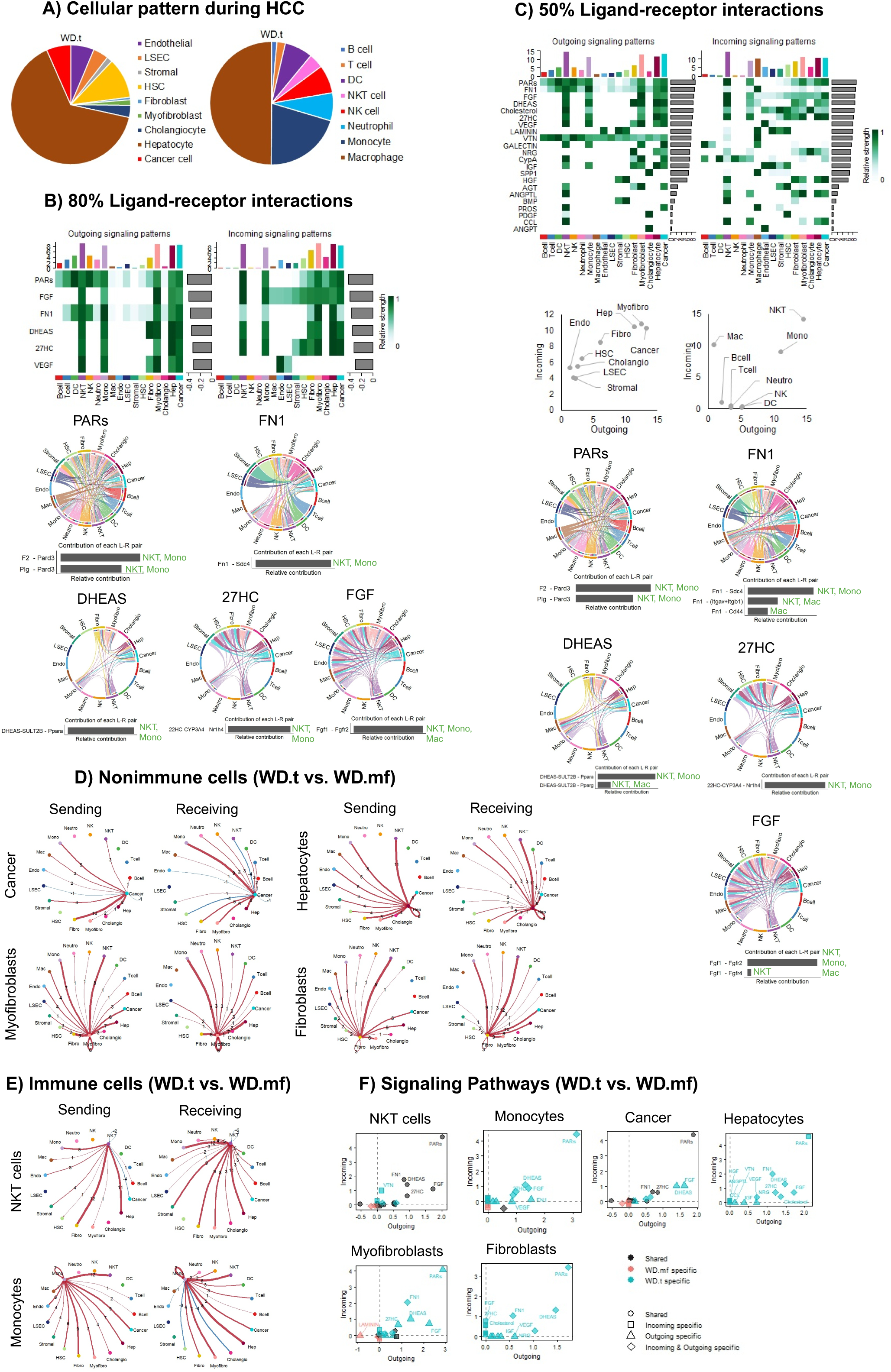
Cancer cells, hepatocytes, and myofibroblasts drive NKT and monocyte-led immune responses via homeostatic pathways in HCC on a WD. A) Immune and nonimmune cell proportions quantified and normalized to 100% in the WD.t group (n=3). B) 80% CellChat analysis heatmap showing all L-R interactions in the WD.t group (upper), and representative chord diagrams showing signaling directionality in PARs, FN1, DHEAS, 27HC, and FGF pathways and their L-R contributions (lower). C) 50% CellChat analysis heatmap portraying all detected L-R pathway interactions in the WD.t group (upper panel), quantified incoming and outgoing interaction strength in nonimmune and immune cell populations (middle panel), and chord diagrams of signaling directionality in PARs, FN1, DHEAS, 27HC, and FGF pathways and their L-R contributions (lower panels). D) Dominant nonimmune cell differential number of interactions sent (left column), and received (right column) in cancer, hepatocytes, myofibroblasts, and fibroblasts in WD.t compared to WD.mf in 50% analyses, showing increased signaling events in red and decreased events in blue in the WD.t group compared to WD.mf, and thicker red or blue arrows show higher increases or lower decreases in signaling events across groups. E) Dominant immune cell differential number of interactions sent (left column) and received (right column) in NKT cells and monocytes in 50% analyses, showing increased number of signaling events in red and decreased in blue for the WD.t group compared to WD.mf, and thicker red or blue arrows show higher increases or lower decreases in signaling events across groups. F) The 50% threshold analysis of signaling pathway changes in cancer, NKT, hepatocyte, myofibroblast, fibroblast, and monocyte cell populations in WD.t compared to WD.mf. Green text in L-R contributions denotes the cell type expressing receptors focused mainly on immune cells, and Itga4+Itgb1 denotes VLA-4.

In order to determine signaling pathways induced in the presence of HCC, comparative analyses between the WD.t vs. WD.mf groups were performed on 50% threshold analysis. These predominant nonimmune cells communicated highest number of signals with monocytes and NKT cells, with myofibroblasts also showing higher number of signaling communications with fibroblasts during HCC (Figure 5D). Compared to NKT cells, monocytes sent highest number of signals to cancer cells (Figure 5E). All these predominant cells upregulated CD1d during HCC (Figure S7), which could present glycolipid to NKT cells for their activation. All other hepatic structural cells communicated predominantly with monocytes or NKT cells (Figure S8A). All other immune cells sent highest number of signals to NKT cells, but they received highest number of signals from monocytes, myofibroblasts or fibroblasts (Figure S8B). Compared to the MAFLD group, the HCC group sent new homeostatic signaling that were also detected in 80% of cells (PARs, FN1, DHEAS, 27HC and FGF) in monocytes, myofibroblasts and fibroblasts, while these signals were relatively increased in NKT cells during HCC (Figure 5F). NKT cells expressed negligible levels of cytotoxic granules whereas between 5-7.5% of T cells and NK cells expressed granzyme A and B as well as 5% of T cells expressing perforin (Figure S9A). Monocytes that were dominating signaling network showed a high level of IL7 expression up to 30% while NKT cells showed a low level up to 20% (Figure S9A). NKT cells lacked cytotoxic function while 5% of T cells targeted 6% of fibroblasts through the TRAIL-TRAILR2 cytotoxic pathway, and 15% of T cells targeted 30% of cancer and 20% of endothelial cells through Fas-mediated cytotoxic pathway (Figure S9B). All the target cells expressed MHC class I (Figure S9C). These data suggest that monocytes become the main immune cells that dominate tissue homeostatic signaling network during HCC.

### Monocyte-driven liver homeostatic pathways can prevent HCC through targeting mainly myofibroblasts, fibroblasts, hepatocytes and cancer following dietary correction

In order to determine whether correction of diet during MAFLD can protect animals against HCC and restore immune cell interaction network similar to that in the CD control group, animals were switched to a CD after 36 weeks of being on a WD. While hepatocytes, cancer and macrophages predominated cellular pattern during tumor progression, monocytes and neutrophils emerged as predominant cells during protection against HCC (Figure 6A). However, analysis of the ligand-receptor interactions representing 80% of cells showed a functional predominance of cancer, stromal cells, cholangiocytes and DCs in sending PARs signaling to themselves but not to hepatocytes during tumor progression (Figure 6B, left panels). On the other hand, protection against HCC was associated with the functional predominance of monocytes, myofibroblasts and fibroblasts in sending the homeostatic signals PARs, DHEAS, NRG and PROS to themselves and to macrophages, hepatocytes and cancer cells (Figure 6B, right panels). Also, all immune cells followed monocytes in PARs signaling pathway (Figure 6B, right panels).

**Figure 6.**
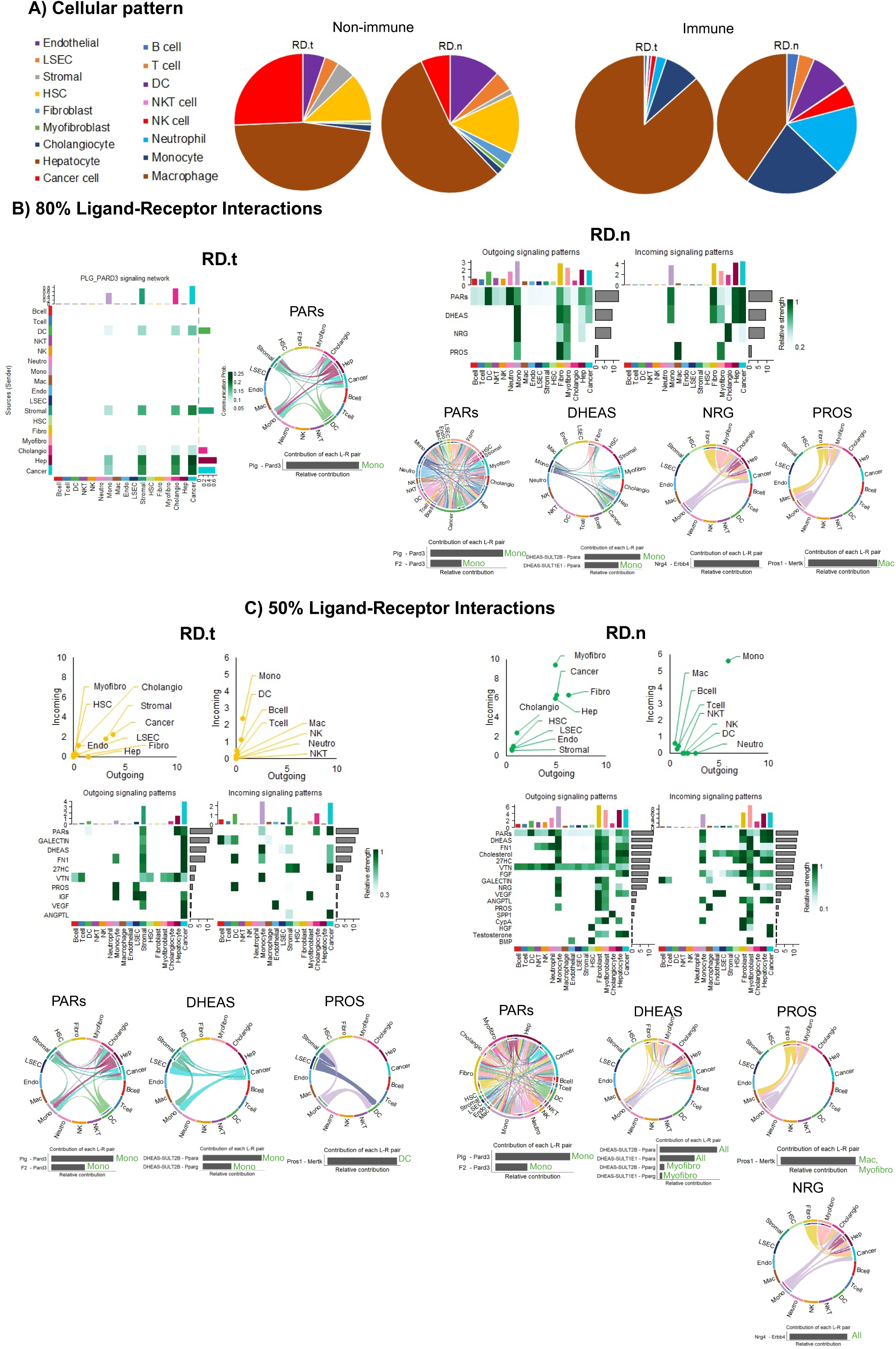
Stromal and cancer cell interactions along with inefficient homeostatic immune responses fuel HCC progression, while myofibroblast and monocyte networks prevent HCC through homeostatic immune pathways after dietary correction. A) Immune (right panel) and nonimmune (left panel) cell proportions quantified and normalized to 100% in the RD.t (n=2) and RD.n (n=3) groups. B) 80% CellChat analysis heatmap in the RD.t and RD.n groups (upper), and the representative chord diagram of PARs in the RD.t group and PARs, DHEAS, NRG, and PROS in the RD.n group signaling directionality and the L-R contributions (lower). C) 50% CellChat analysis quantified incoming and outgoing interaction strength for nonimmune and immune cell populations (upper panel), heatmap of detected L-R pathways (middle panel), and chord diagrams of signaling directionality for PARs, DHEAS, PROS and NRG pathways and their L-R contributions (lower panel) in the RD.t (left) and RD.n (right) groups. Green text in L-R contributions denotes the cell type expressing receptors and focused mainly on immune cells and Itga4+Itgb1 denotes VLA-4. Green text in L-R contributions denotes the cell type expressing receptors and focused mainly on immune cells and Itga4+Itgb1 denotes VLA-4.

For 50% of cells, stromal cells and cancer cells were predominantly involved in sending the homeostatic signals (PARs, DHEAS, 27HC, VTN, ANGPTL) to themselves or monocytes as well as sending VEGF to endothelial cells during tumor progression (Figure 6C, left panels, Figure S10). In this group, monocytes produced FN1 and PROS for targeting cancer and DCs as well as IGF for targeting myofibroblasts (Figure 6C, left panels, Figure S10). On the other hand, protection against HCC was associated with monocyte-dominated homeostatic signals (FN1, PROS, PARS, DHEAS, Cholesterol, 27HC, VTN and NRG) that targeted mainly myofibroblasts, fibroblasts, hepatocytes and cancer cells, as well as VEGF that targeted endothelial cells and LSEC (Figure 6C, right panels, Figure S10). Myofibroblasts and fibroblasts produced additional homeostatic molecules (ANGPTL, CypA, HGF and Testosterone) that targeted themselves, monocytes, hepatocytes or cancer (Figure 6C, right panels, Figure S10). DHEAS was reported to induce cell death in liver tumor cells ^51,52^. It will inhibit myofibroblast activation and reducing ECM production as well as induce apoptosis in myofibroblasts. These prevent liver stiffening associated with chronic fibrosis and HCC. The PROS1-MERTK axis is crucial for efferocytosis by macrophages, which helps resolve inflammation and apoptotic cells. Also, NRG4 might influence cholangiocyte proliferation, potentially aiding in maintaining bile duct integrity during hepatic stress. as well as cholangiocytes survival through Erbb4 signaling resulting in the suppression of NASH-HCC ^53^.

### Prevention of HCC is associated with elevated homeostatic pathways that are distinct from the CD group

In order to determine whether protection against HCC following dietary correction was associated with the restoration of the signaling network as in the healthy CD group, comparative analysis of dominant cellular interaction signaling was performed at the 50% threshold setting. Compared to the CD group, monocytes sent the highest number of signals to stromal and cancer cells during HCC (RD.t) whereas the highest signals targeted myofibroblasts and fibroblasts during the inhibition of HCC (RD.n) (Figure 7A). Myofibroblasts and fibroblasts sent highest number of homeostatic signals to themselves only during tumor inhibition in the RD.n group (Figure 7A). Stromal cells and cancer cells sent the highest number of homeostatic signals to themselves and monocytes during tumor progression whereas they targeted mainly myofibroblasts during tumor inhibition (Figure S11). Such distinct cellular targeting was also observed for all other hepatic structural cells (Figure S11). All other immune cells of animals protected against HCC following dietary correction showed the highest number of homeostatic signaling communications mainly to myofibroblasts and fibroblasts whereas such signals decreased during HCC progression of a RD when compared to the CD group (Figure S12). Homeostatic signaling involved in anti-fibrotic (Testosterone, HGF) or pro-fibrotic functions (FGF, Cholesterol, CypA, Spp1) were detected only during protection against HCC (Figure 7B). On the other hand, fibrinogenic IGF signaling targeting myofibroblasts was detected only during HCC (Figure 7B). Four pathways (27HC, VTN, FN1, ANGPTL) that were active in both groups were found to target mainly monocytes and myofibroblasts only in the RD.n group (Figure 7B). The Itgb5 axis in myofibroblasts regulates ECM turnover and facilitates liver regeneration after injury on myofibroblasts are tightly regulated, preventing excessive ECM deposition or prolonged myofibroblast activation. The Itgb1 axis allows myofibroblasts to sense and respond to mechanical changes in the liver microenvironment, regulate ECM dynamics and maintain tissue homeostasis. This balance ensures tissue integrity without progressing to fibrosis ^54^. The CD44 pathway promotes ECM remodeling, hyaluronan clearance, and transient activation for wound healing. The SDC4 stabilizes cell-ECM interactions, coordinates ECM turnover, and the SDC2 pathway activates RhoA/ROCK, promoting cytoskeletal reorganization, and promote myofibroblasts survival by activating PI3K/Akt and ERK/MAPK pathways. In monocytes, the SDC4 pathway supports monocyte differentiation into macrophages with enhanced phagocytic activity, crucial for clearing apoptotic cells and debris but also contributing to fibrosis under chronic conditions.

**Figure 7:**
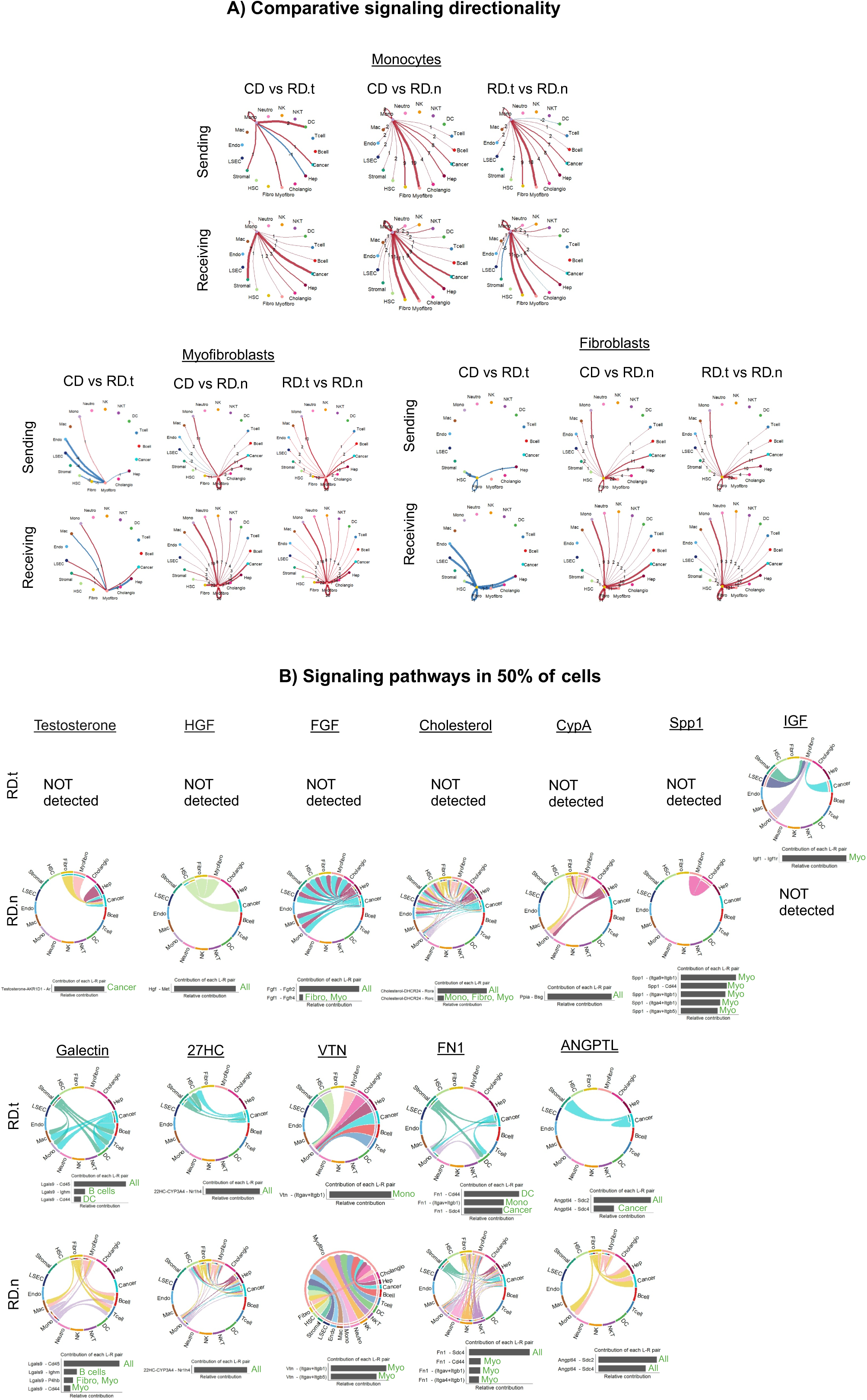
Hepatic homeostatic pathways associated with immune responses during the diet reversal-induced protection against HCC. A) Differential number of interactions sent (upper rows), and received (lower rows) in Monocytes, myofibroblasts, and fibroblasts in RD.t compared to CD (CD vs RD.t; left columns), RD.n compared to CD (CD vs RD.n; middle columns), and RD.n compared to RD.t (RD.t vs RD.n; right columns) in 50% analyses; showing increased signaling events in red and decreased events in blue for RD.t compared to CD, RD.n compared to CD, and RD.n compared to RD.t; thicker red or blue arrows show higher increases or lower decreases in signaling events across comparisons. B) Chord diagrams depicting signaling directionality of pathways detected in 50% CellChat analyses for RD.t (upper row) and RD.n (lower row) groups. Ligand-receptor (LR) contributions are shown below, with green text indicating the cell types receiving signals in 50% threshold signaling pathways from the chord diagrams.

Cytotoxic immune responses were detected only in 5-10% of NK cells and T cells (Figure S13A). While expression of a crucial homeostatic cytokine IL7 that plays a vital role in the survival and maintenance of immune cells was negligible in the RD.t group, up to 30% of monocytes expressed IL7 in the RD.n group (Figure S13A). Also, high level of CXCL13 was expressed by 80% of monocytes and macrophages, and high level of CCL4 was expressed by 15% of monocytes and macrophages in the RD.t group (Figure S13A). Elevated CXCL13 levels are associated with poor prognosis in HCC ^55^. It fosters a tumor-promoting microenvironment by recruiting regulatory B cells and T cells that suppress anti-tumor immune responses ^56^. Also, CCL4 has been shown to be involved in liver injury, enhancing lipotoxic stress in hepatocytes ^57^. All hepatic cells expressed MHC class I (Figure S13B). Finally, 5-10% of T cells expressed high levels of Fas-L that targeted 20-30% of myofibroblasts, fibroblasts and endothelial cells turnover mainly in the RD.n group (Figure 13C).

### Catalase-expressing monocytes may repair the tumor microenvironment

Since monocytes were found to dominate the receptor-ligand cell-cell network during HCC (WD.t) and protected against HCC (RD.n) with a reduced functional interactions in the HCC on a reverse diet (RD.t), we analyzed antioxidant pathways and scavenger pathways through which entropy of the TME is reduced by monocytes. We found that 60% of monocytes expressed high levels of catalase (Cat) in the WD.t and RD.n groups only (Figure 8A) while 40% of monocytes in the RD.t group expressed high levels or MerTK (Figure 8B). MerTK is a receptor tyrosine kinase that binds to apoptotic cells via ligands like Gas6 and Protein S, which recognize phosphatidylserine on the surface of dying cells, resulting in efferocytosis of dead cells. Catalase is an important antioxidant enzyme that reduces oxidative stress by breaking down hydrogen peroxide (H□O□) into water and oxygen, and reduces ROS levels. Similarly, the PARs pathway (Plg-Pard3) mainly by monocytes and cancer cells targeted hepatocyte regeneration as well as liver structural cells for ECM remodeling in the WD.t and RD.n groups (Figure 8C-D). However, the PARS signaling was shifted to cancer and stromal cells targeting themselves during the progression of HCC after dietary correction (Figure 8E). These data suggest that monocyte-driven catalase and PARS pathways that detoxify the liver and induce liver regeneration, respectively, prevents HCC unless they are overwhelmed by daily intake of carcinogenic WD such that retention of these pathways following dietary correction resulted in tumor inhibition whereas loss of the liver regeneration pathway led to HCC progression.

**Figure 8.**
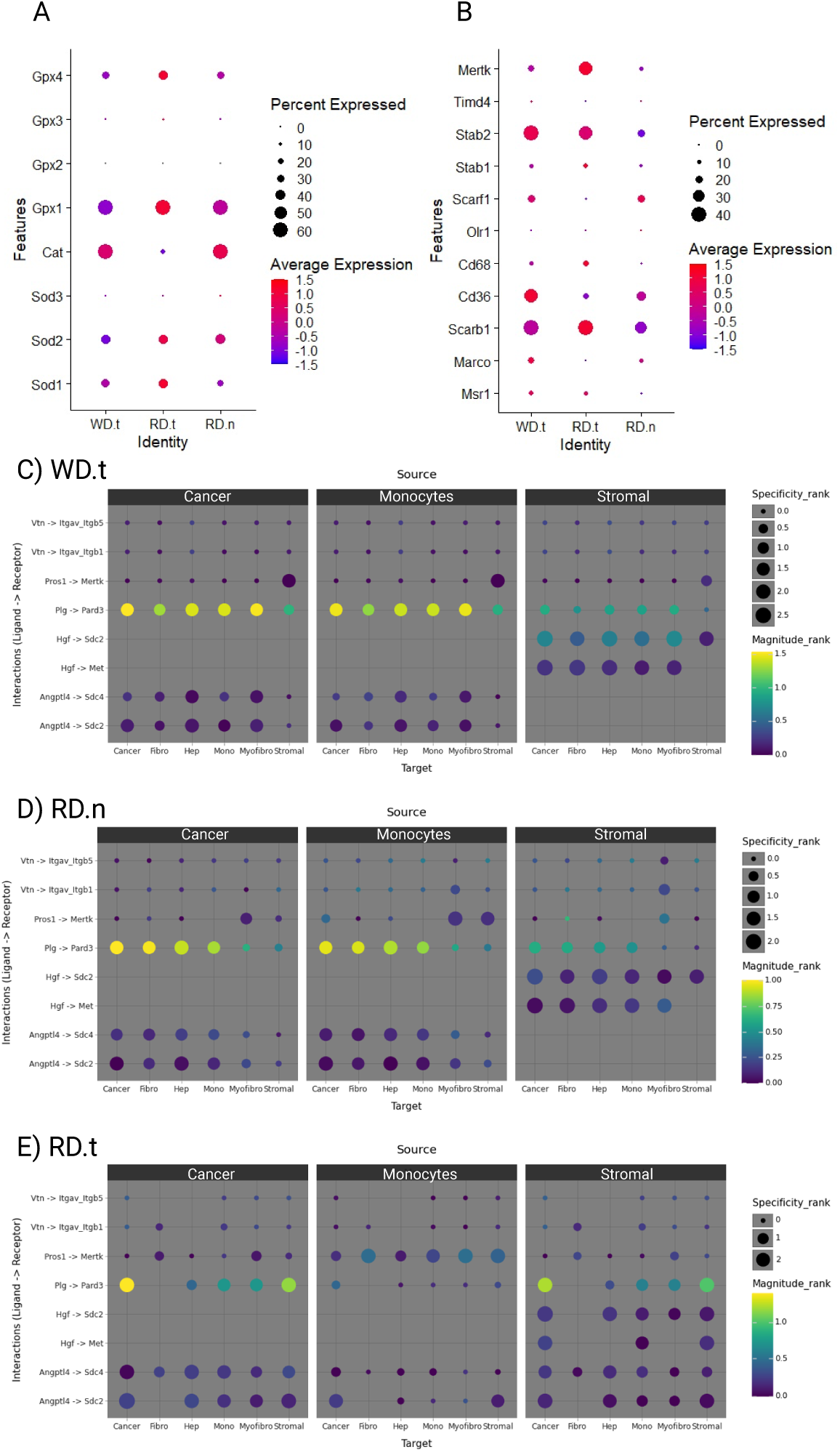
Monocytes function during HCC progression or inhibition. DotPlots showing percentage of monocyte population expression and log-normalized average transcript expression for antioxidant enzymes (A) and scavenger receptors (B) during HCC on a WD (WD.t), HCC on a reverse diet (RD.t) and protected against HCC following diet reversal (RD.n). C-E) A <u>LI</u>gand-receptor <u>AN</u>alysis fr<u>A</u>mework (LIANA) was used to confirm key interactions detected in the HCC tumor-bearing mice (C), and mice that underwent diet reversal and did not show HCC tumors (D) and those that did (E). The LIANA Rank Aggregate method was applied to each dataset individually and represents a consensus by integrating the predictions of individual methods from multiple programs LR detection methods (CellPhoneDB, Connectome, log2FC, NATMI, SingleCellSignalR, Geometric Mean, scSeqComm, and CellChat). The results were filtered to show LR interactions detected in 50% CellChat analyses that were also present in the LIANA consensus database (Plg-Pard3, Pros1-Mertk, Vtn-Itgav_Itgb1, Vtn-Itgav_Itgb5, Angptl4-Sdc2, Angptl4-Sdc4, and Hgf-Met). Specificity rank shows how specific an interaction is to a pair of cell types, in which larger the dot size, the more specific to a pair of cell types it is. Magnitude rank shows the corresponding strength of that detected LR interaction, with yellow being stronger and blue being weaker.

## Discussion

Our previous research demonstrated cancer cell dormancy in the FVBN202 transgenic mouse model of spontaneous breast cancer ^25^, as well as hepatic carcinogenesis events in the DIAMOND model of MAFLD progression to HCC ^9^. In this study, we employed a systems immunology approach to dissect how immune cells interact within complex cellular networks to coordinate hepatic immune responses. Our findings reveal that carcinogenesis begins during MAFLD by detecting the oncogenes GPC3 and AFP, but remain dormant because of the highest level of the Hnf4a tumor suppressor gene whereas Hnf4α dropped during the progression of MAFLD to HCC.

Contrary to the long-held view of the liver as a tolerogenic organ, we show that the liver actively engages immune responses predominantly aimed at maintaining tissue homeostasis. Under healthy condition, up to 25% of immune cells become activated by hepatic structural cells (Figure 2D) and up to 50% of macrophages received GAS signaling from cholangiocytes to promote efferocytosis (Figure 2C-D). During a WD-induced MAFLD, up to 80% of immune cells received activating signals (Gal9, PARS, FN1) and inhibitory signal (Cholesterol) with PARS and FN1 continue to activate immune cells but with the addition of more inhibitory signals (DHEAS, FGF) during HCC (Figure S10). Cholesterol has been shown to induce CD8^+^ T cell exhaustion in the TME ^58^, disrupt T cell activation ^59^, and facilitate tumor immune evasion ^60^. Dietary correction shifts NKT cell-dominated pattern to monocyte-dominated homeostatic responses. Importantly, the efficacy of these monocyte-driven responses hinges on their ability to target myofibroblasts and fibroblasts, crucial for restoring liver tissue integrity and preventing HCC. Inadequate monocyte targeting of these cells leads to failure in tissue repair and progression to HCC (Figure S10).

Our data challenges the notion of liver being a tolerogenic organ by demonstrating immune cell activation networks in the liver during health and disease. Such tissue homeostatic immune response has been demonstrated by the presence of tissue-resident T memory (Trm) cells intrinsically tied to tissue homeostasis ^61^. In fact, Trm cells demonstrate tissue remodeling abilities by producing the epidermal growth factor receptor (EGFR)-ligand amphiregulin (AREG), which promotes epithelial cell regeneration. Notably, blocking EGFR signaling or the cytokines IFN-γ and TNF has been shown to inhibit this tissue remodeling process ^62^. These findings align with reports of hepatic CXCR6+ NKT cells inhibiting MAFLD progression to HCC ^63^ and the role of therapies like Resmetirom in restoring liver homeostasis ^64^. Additionally, the host-protective role of NKT cells has been documented in mouse models induced by fast food diet (FFD) and methionine-choline-deficient (MCD) diet; however, chronic administration of α-GalCer can induce NKT cell anergy and promote disease progression ^40^, suggesting that NKT cells may become overwhelmed by chronic WD intake. Finally, monocyte-dominated network in the WD.t and RD.t group was associated with increased catalase that neutralizes ROS, thereby protecting immune cells from oxidative damage, ensuring their proper function in the TME. Given the key role of ROS in the pathogenesis of MAFLD and HCC, such antioxidant function of monocytes can be a key contributor to the TME remodeling, though in the WD.t group, it could be overwhelmed by chronic intake of WD. To this end, excess dietary fat leads to the accumulation of free fatty acids (FFAs) in hepatocytes, which undergo β-oxidation in mitochondria, producing ROS as byproducts. Also, fructose metabolism increases substrate overload for mitochondrial oxidative phosphorylation, generating excessive ROS. However, diet reversal facilitates the function of catalase to repair the liver in the RD.n group while its absence in the RD.t group promotes HCC. Similar trend was detected for monocyte-driven PARS pathway targeting liver regeneration to be inhospitable for the tumor while stromal- and cancer-driven PARS pathway in an autocrine pathway facilitated HCC progression even during dietary correction. Therefore, different cellular targets of PARS pathways result in distinct outcomes. Our findings align with previous reports highlighting the critical role of the monocyte-target axis^65^ and catalase^66^ in mediating dual functions that either promote or inhibit tumor progression. Monocyte-driven immune regulation, as illustrated through source-target cell communication pathways, can be summarized in a model (Figure S10) that underscores key homeostatic pathways driving HCC progression or inhibition following dietary correction. This model warrants further validation through future proteomic studies.

This study challenges the traditional paradigm of tumor immunology, demonstrating that hepatic immune responses during MAFLD and HCC are primarily homeostatic rather than cytotoxic. This concept is supported by other reports showing the role of immune cells in liver homeostasis ^4,5,62,67^. Although commercially available antibodies for detecting immune networks limit the ability to verify discovered pathways at the protein level, further validation can be achieved using proteomic approach in future studies. Alterations in hepatic homeostasis driven by WD not only initiate carcinogenesis but also activate protective immune mechanisms through enhanced homeostatic pathways. Dietary correction alleviates carcinogenic pressures, enabling monocyte-driven responses to restore tissue integrity and prevent HCC. These insights redefine immunotherapy strategies for MAFLD and HCC, advocating for interventions that enhance immune-mediated tissue repair and homeostasis, rather than focusing exclusively on direct cytotoxicity. This paradigm shift opens the door for innovative treatments that leverage the immune system’s natural ability to maintain tissue health and integrity, potentially revolutionizing cancer prevention and therapy.

## Authors’ Contributions

Conceptualization, M.H.M.; investigation, N.K., F.M., H.F.A., M.S., M.O.I.; writing first draft: N.K., M.H.M.; all authors edited the manuscript and approved the final submission; supervision: F.M., A.L.O., A.J.S, and M.H.M.; funding acquisition, M.H.M. and A.J.S. All authors reviewed the manuscript.

