## Supplemental data for "Anti-tumor immunity relies on targeting tissue homeostasis through monocyte-driven responses rather than direct tumor cytotoxicity"

A) Experimental Design

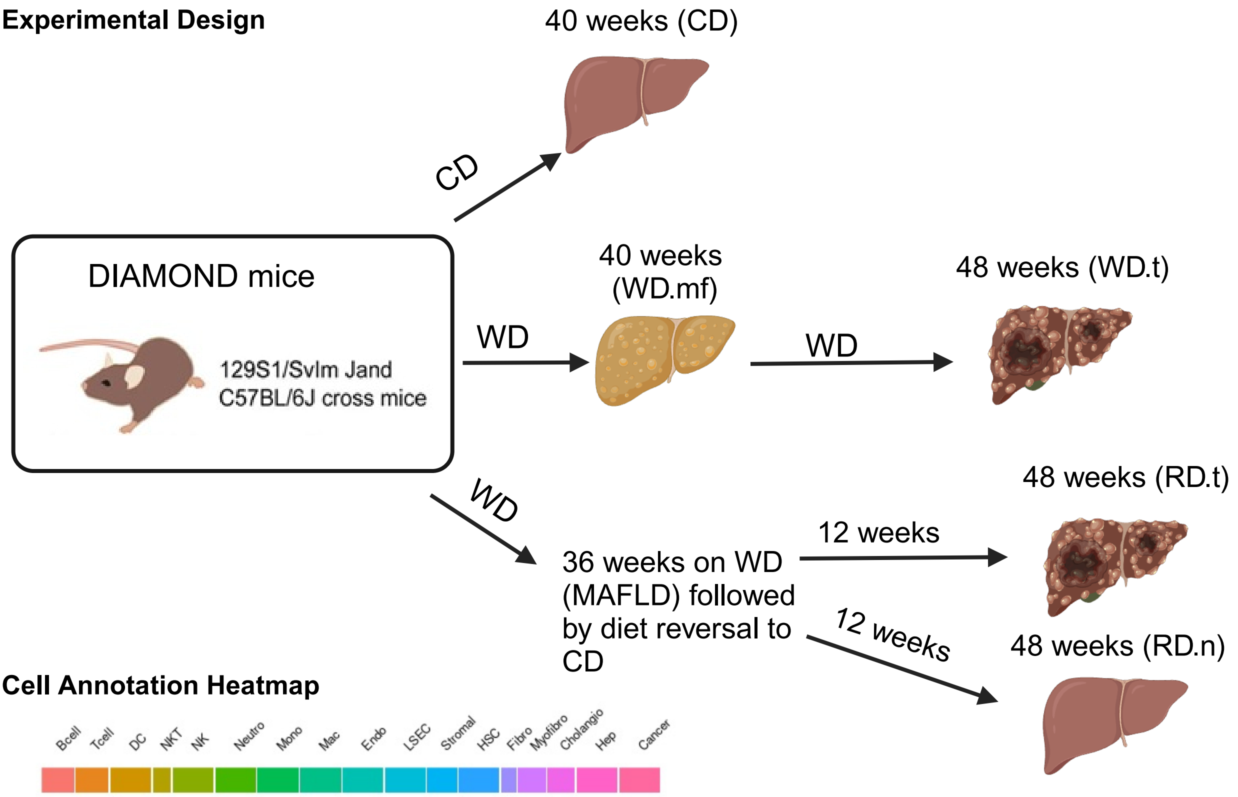


B) Cell Annotation Heatmap

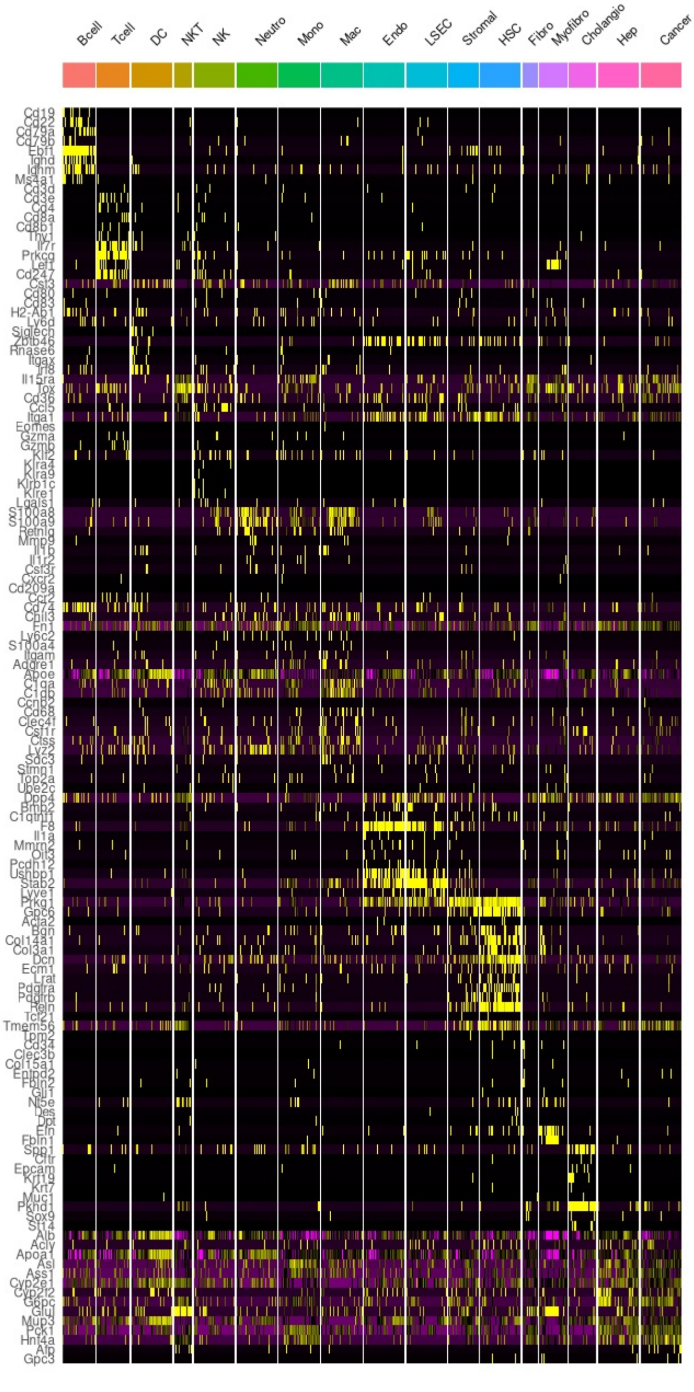


**Figure S1. Experimental design and Cellular Annotation.** A) Diet-induced animal model of nonalcoholic fatty liver disease (DIAMOND) mice were put on a standard chow diet (CD) for 40 weeks, WD for 40 weeks that developed MAFLD (WD.MAFLD, WD.mf), or a WD continued to 48 weeks that developed HCC (WD.HCC, WD.t). In addition, mice were on a WD for 36 weeks and underwent diet reversal back to a CD and followed up to 48 weeks of age, which some mice developed HCC (RD.HCC, RD.t) and others were protected (RD. no HCC, RD.n). B) Heatmap showing the marker genes from the CellMarker2.0 database utilized through scSorter for identifying and annotating cell types within the sequencing samples.

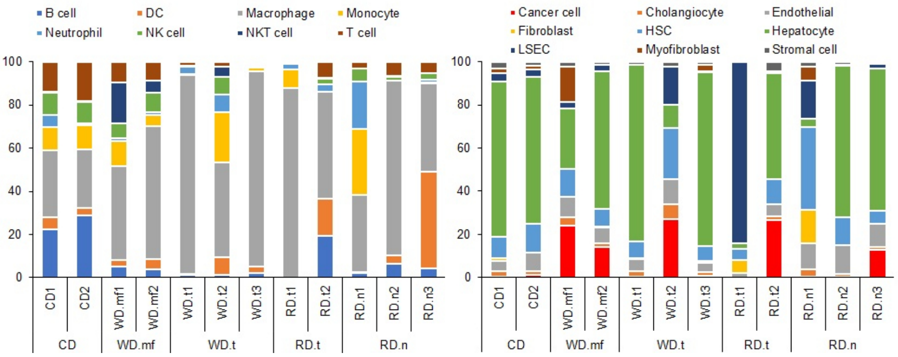


**Figure S2. Individuality of snRNA-seq sample cell annotations.** Bar graphs showing cell annotations from scSorter quantified and normalized to 100% for individual samples for immune cells (left) and non-immune cells (right) in the chow diet group (CD; n=2), metabolic-associated fatty liver disease group (MAFLD; n=2) after 40 weeks on western diet (WD.mf), WD-induced HCC tumor bearing mice after 48-52 weeks on WD (WD.t; n=3), and mice that underwent diet reversal back to a CD after 36 weeks of being on WD that showed HCC tumors in the liver (RD.t; n=2) and those without tumors (RD.n; n=3)

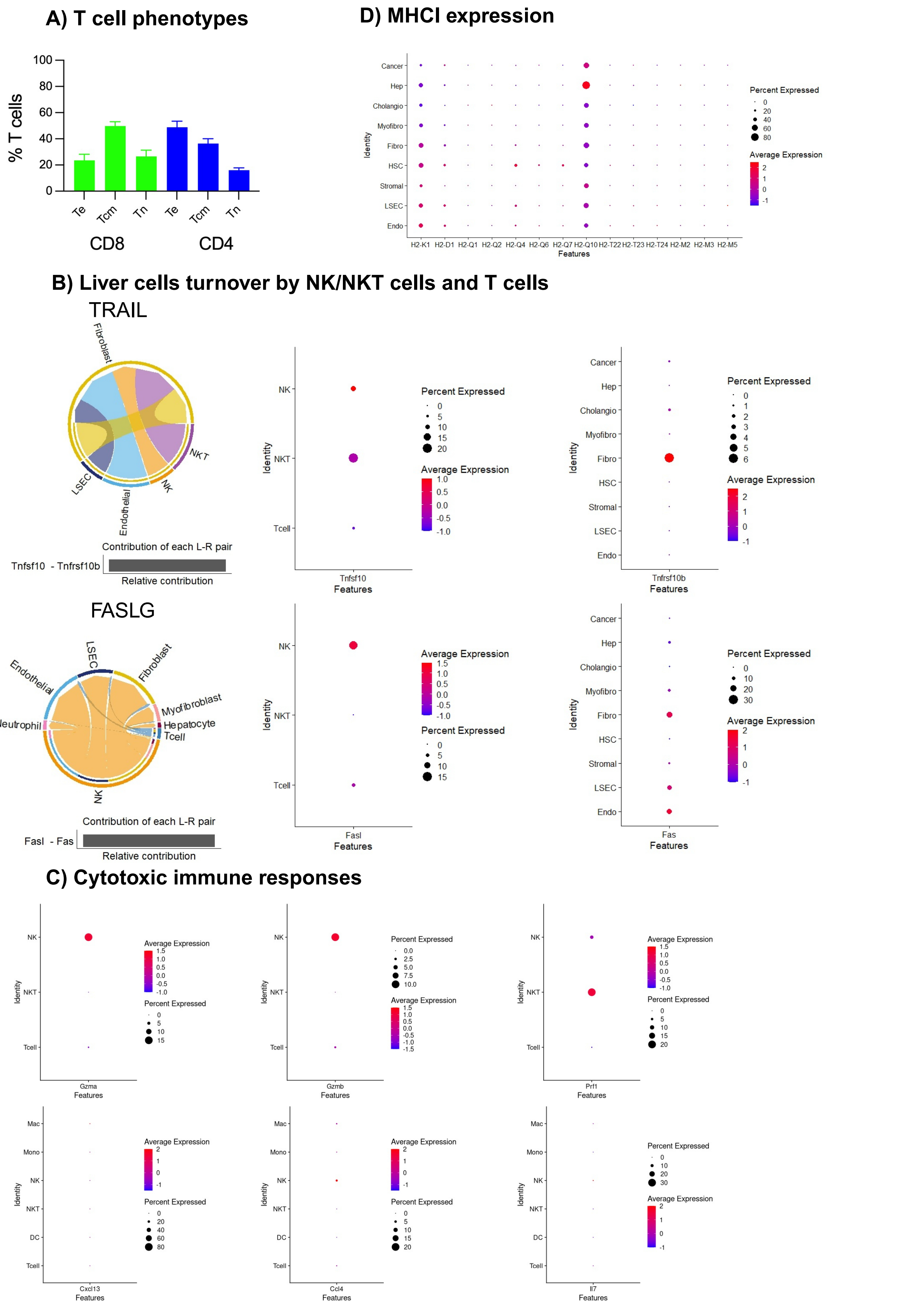


**Figure S3**. A) Detection of hepatic T cell subsets with flow cytometry to differentiate T effector cells (Te, CD44$^+$CD62L$^{-/low}$), T central memory cells (Tcm, CD44$^+$CD62L$^{high}$), and T naive cells (Tn, CD44$^-$CD62L$^+$) in the CD4$^+$ and CD8$^+$ populations. B) Chord diagrams depicting signaling directionality detected in 5% threshold analyses for TRAIL (upper left) and FasLG (lower left) pathways and their L-R contributions, as well as quantified DotPlots showing percent cell population expression and log-normalized average transcript expression of Tnfsf10/Tnfrsf10b (upper right) and FasL/Fas (lower right) in the CD group (n=2). C) DotPlots showing percent cell population expression and log-normalized average transcript expression of Gzma, Gzmb, and Prf1 in NK, NKT, and T cells (upper row), and Cxcl13, Ccl4, and IL7 in NK, NKT, T cell, Macrophages (Mac), Monocytes (Mono), and Dendritic cells (DC) in the CD group (lower row). D) DotPlot showing percent cell population expression and log-normalized average transcript expression of MHCI molecules in structural cells in the CD group.

#### 50% homeostatic pathways

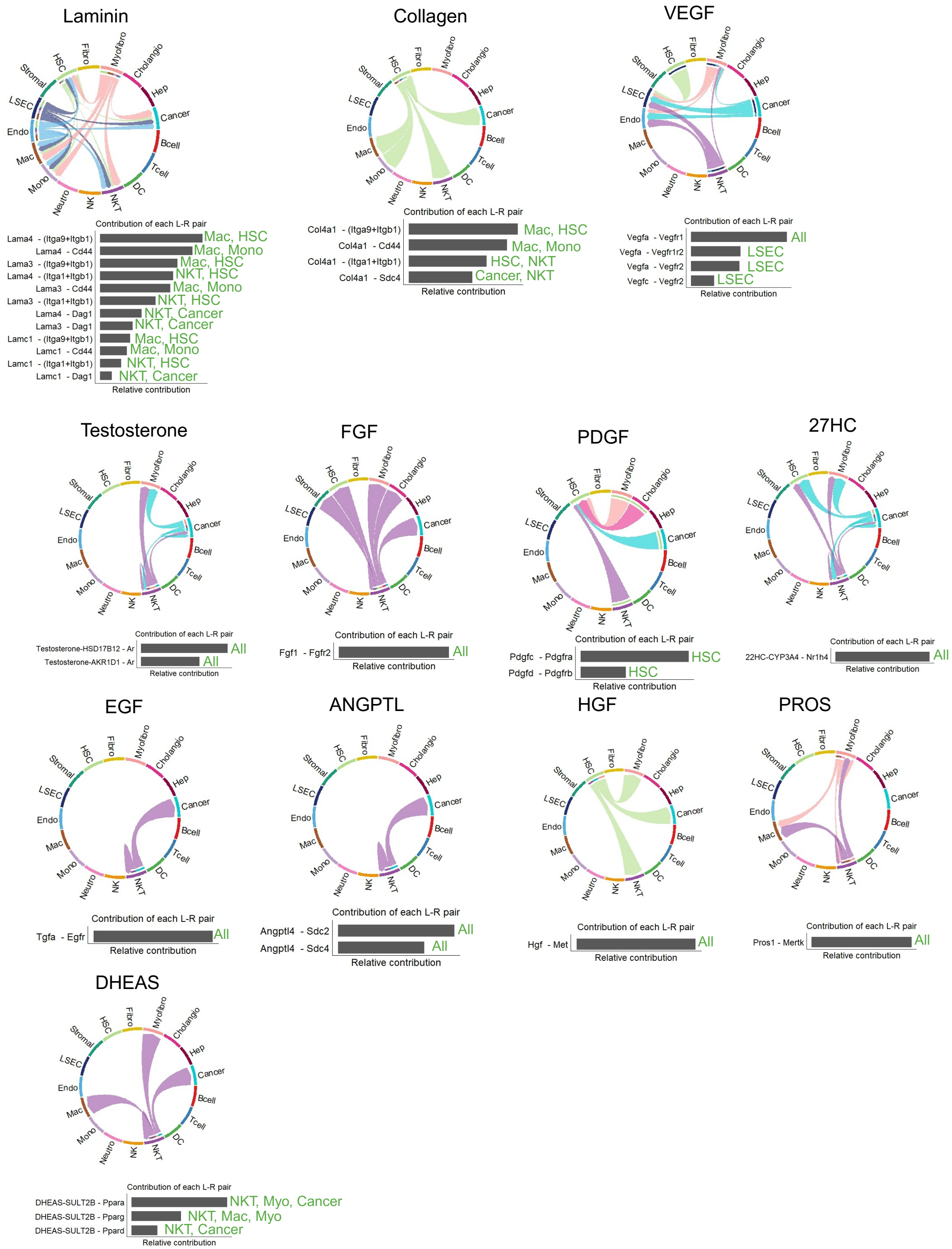


**Figure S4**. Chord diagrams showing signaling directionality of Laminin, Collagen, VEGF, Testosterone, FGF, PDGF, 27HC, EGF, ANGPTL, HGF, PROS, and DHEAS pathways and the L-R contributions (below) in 50% of cells in the WD.mf group (n=2). Green text shows the cell types expressing receptors.

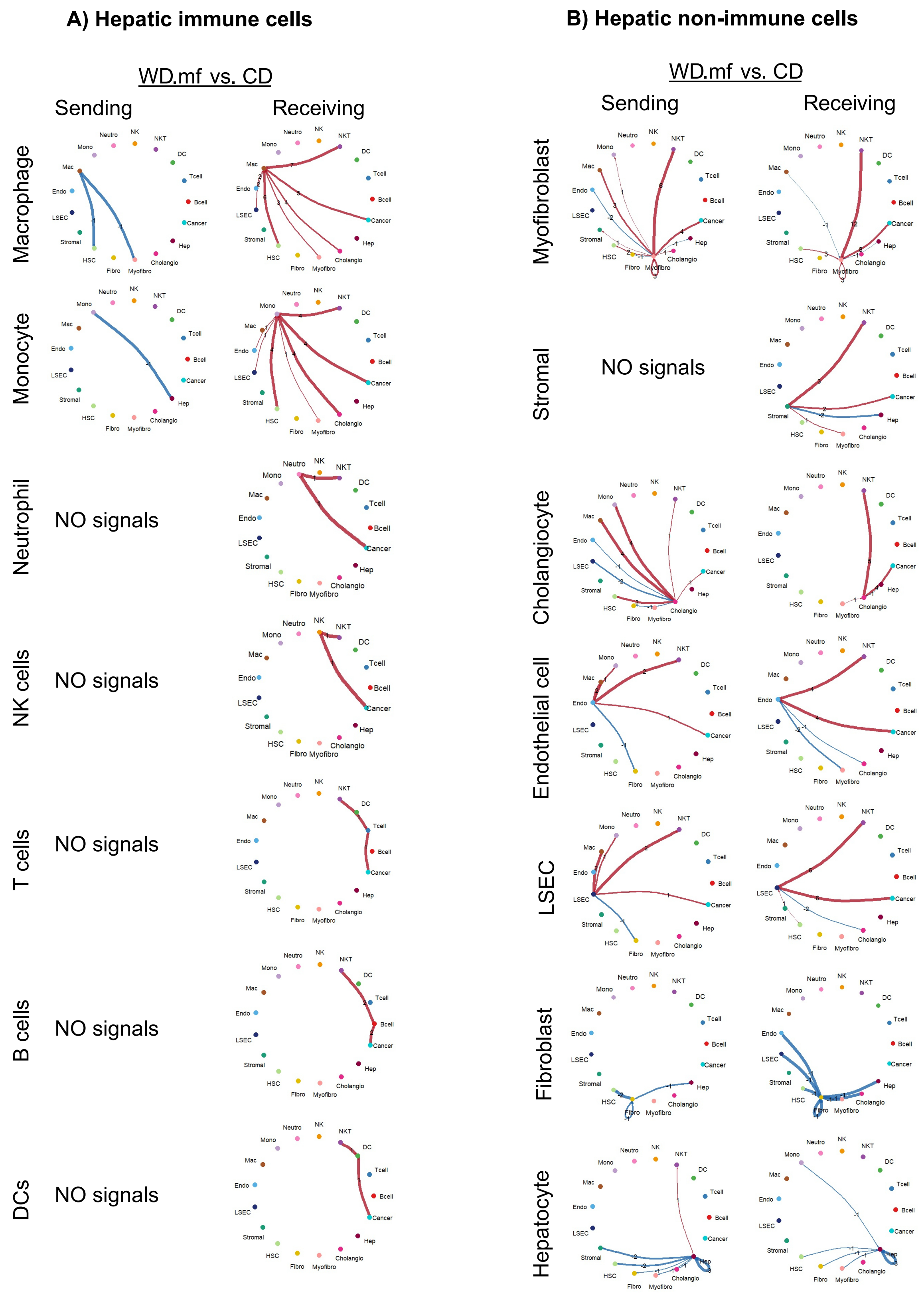


**Figure S5**. A-B) Differential number of signaling interactions in WD.mf compared to CD in 50% threshold analyses sent (left column) and received (right column) for hepatic immune cells (A) and non-immune cells (B); red color shows number of increased interactions and blue shows decreased interactions in the WD.mf group compared to CD group.

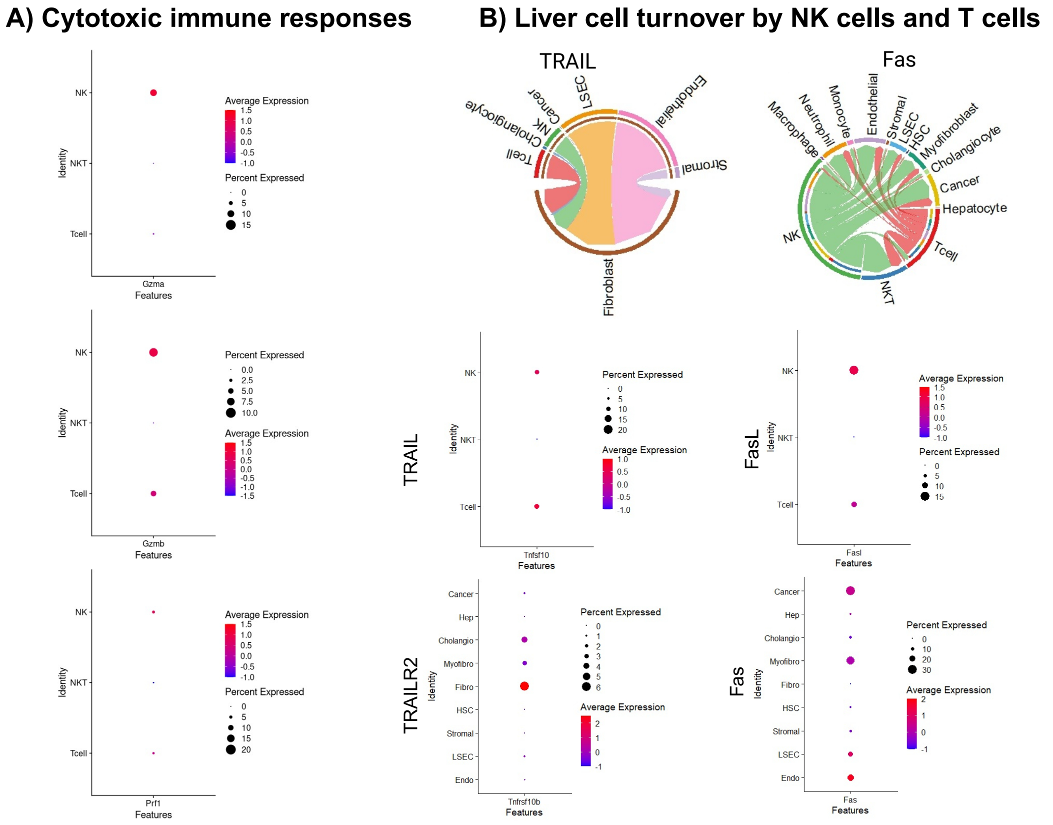


**Figure S6**. A) DotPlots showing percent cell population expression and log-normalized average transcript expression of Gzma, Gzmb, and Prf1 in NK, NKT, and T cells. B) Chord diagrams depicting signaling directionality detected in 5% threshold analyses for cytotoxic pathways (TRAIL and Fas) in the WD.mf group, and DotPlots showing the percentage of expression and log-normalized transcript expression of TRAIL (Tnfsf10), TRAILR2 (Tnfrsf10b), FasL (Fasl), and Fas in the WD.mf group.

#### Cd1d1 Expression

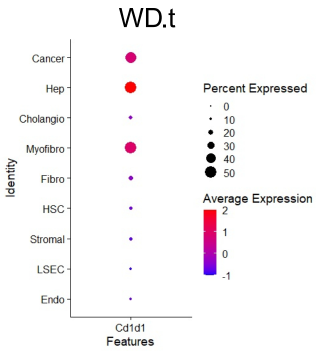


**Figure S7**. DotPlot showing percentage of cell population expression and log-normalized average transcript expression of Cd1d1 in WD.t group structural cells.

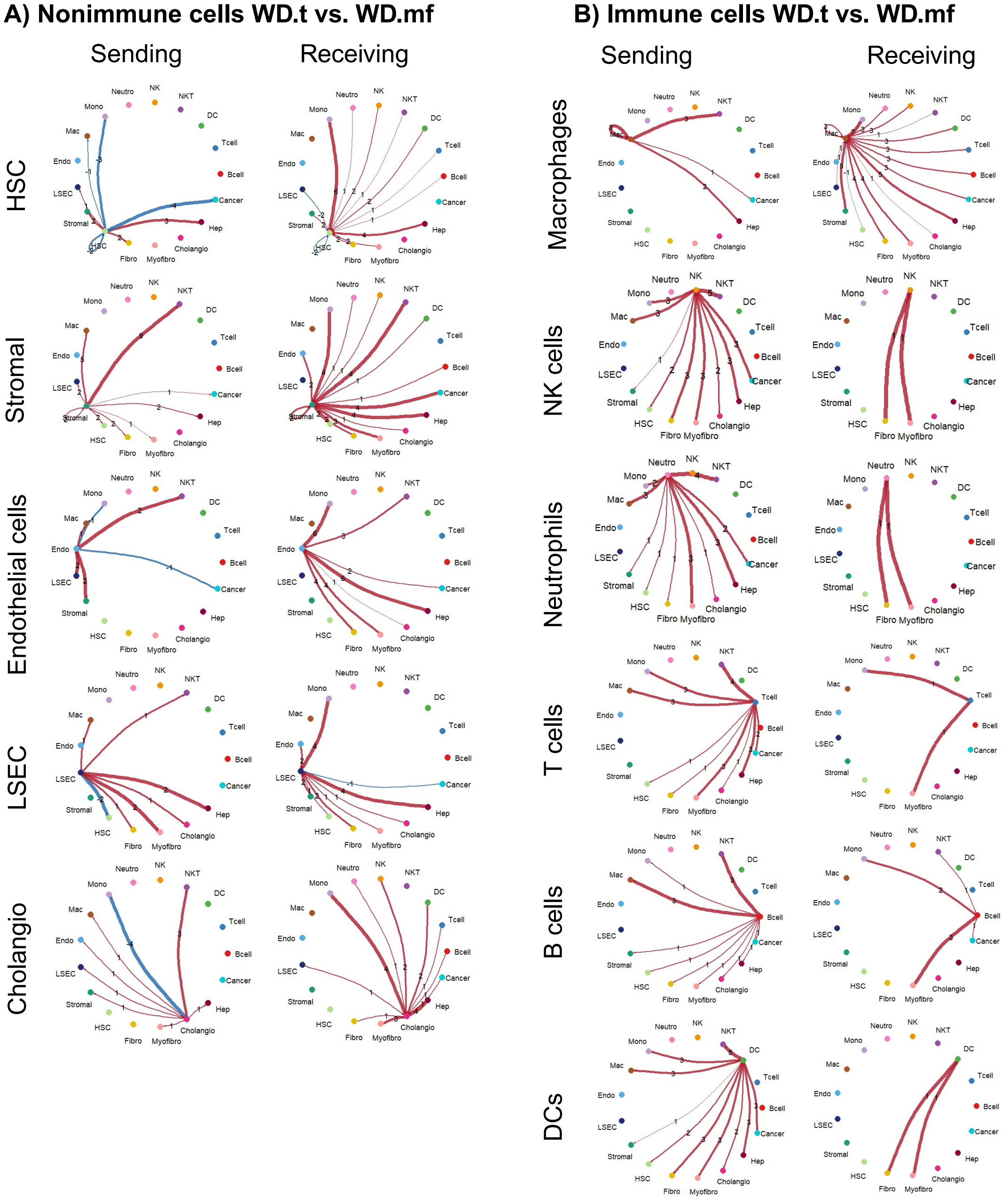


**Figure S8**. A-B) Differential number of signaling interactions detected in 50% threshold analyses sent (left column) and received (right column) in hepatic non-immune cells (A) and immune cells (B) in the WD.t group compared to WD.mf. Red colored arrows show increased number of signaling events and blue colored arrows show decreased events in the WD.t group compared to WD.mf.

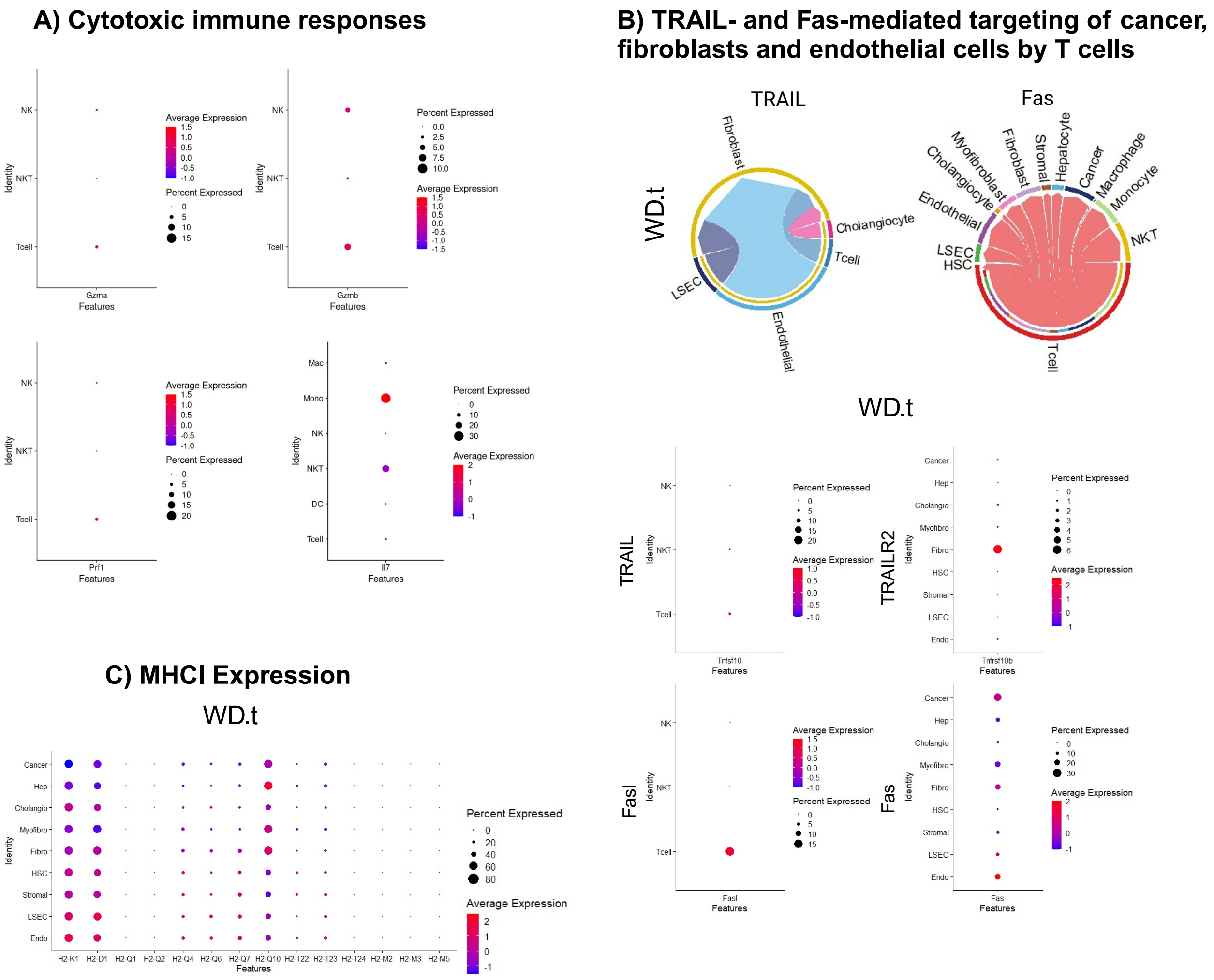


**Figure S9**. A) DotPlots showing percentage of cell population expression and log-normalized average transcript expression of Gzma, Gzmb, Prf1, IL7 in immune cells. B) Chord diagrams depicting signaling directionality detected in 5% threshold analyses for cytotoxic pathways (TRAIL and Fas) in the WD.t group (upper panels). DotPlot showing percentage of cell population expression and log-normalized transcript expression of TRAIL (Tnfsf10, upper left), TRAILR2 (Tnfrsf10b, upper right), FasL (Fasl, lower left), and Fas (lower right) in the WD.t group. C) DotPlot showing percentage of cell population expression and log-normalized average transcript expression of MHCI molecules in the WD.t group structural cells.

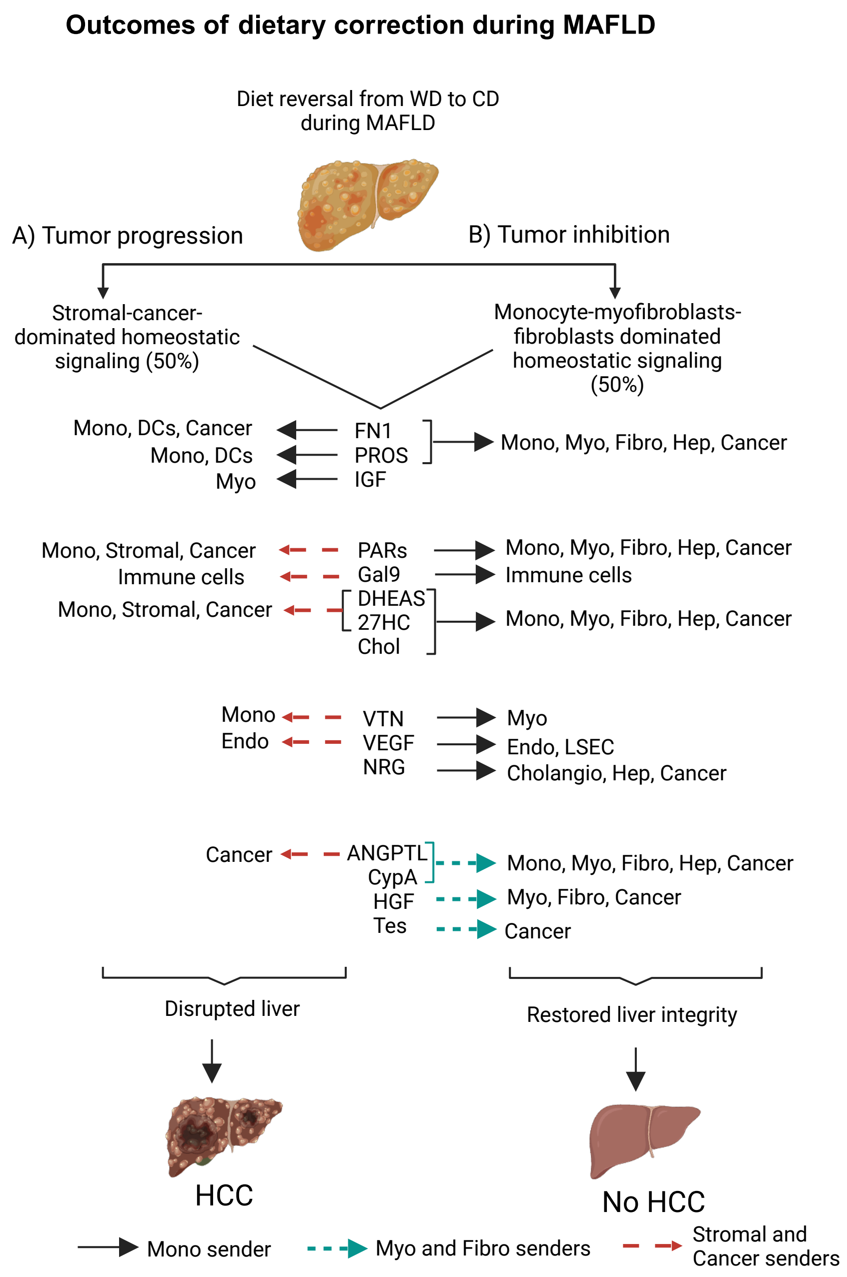


**Figure S10. Monocyte-dominated immune responses restores liver tissue integrity through homeostatic pathways that targets myofibroblasts, fibroblasts, hepatocytes and cancer, thereby preventing HCC.** Diamond mice were switched to a CD during MAFLD and their liver tissue was analyzed 12 weeks after being on a dietary correction. A) The stromal cell-cancer cell-dominated homeostatic pathways targeting mainly stromal cells and cancer promoted HCC whereas B) monocyte-dominated homeostatic pathways targeting myofibroblasts, fibroblasts, hepatocytes and cancer inhibited HCC. There were also additional tissue homeostatic pathways (Cholesterol, NRG, CypA, HGF and Testosterone) present during tumor inhibition while they were absent during tumor formation.

### Signaling directionality in 50% of nonimmune cells

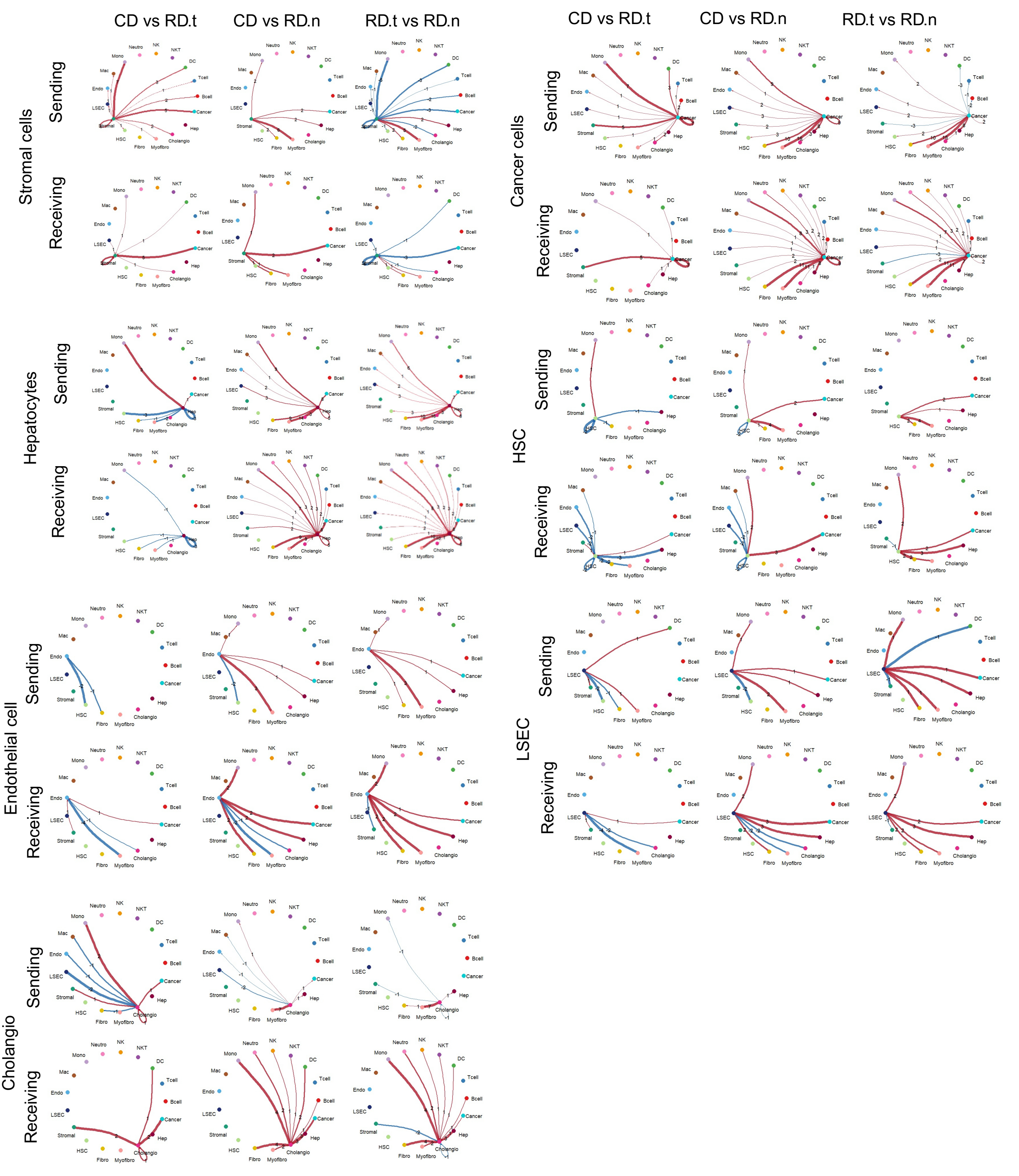


**Figure S11**. The 50% threshold analysis showing the differential number of interactions sent (upper rows) and received (lower rows) for stromal cells, cancer cells, hepatocytes, HSCs, endothelial cells, LSEC and cholangiocytes (Cholangio) across multiple comparisons such as CDvRD.t (left columns), CDvRD.n (middle columns), and RD.tvRD.n (right columns). Red arrows depict increased number of signaling events and blue arrows show decreased events in the RD.t group compared to CD (CDvRD.t), RD.n group compared to CD (CDvRD.n), and RD.n group compared to RD.t (RD.tvRD.n).

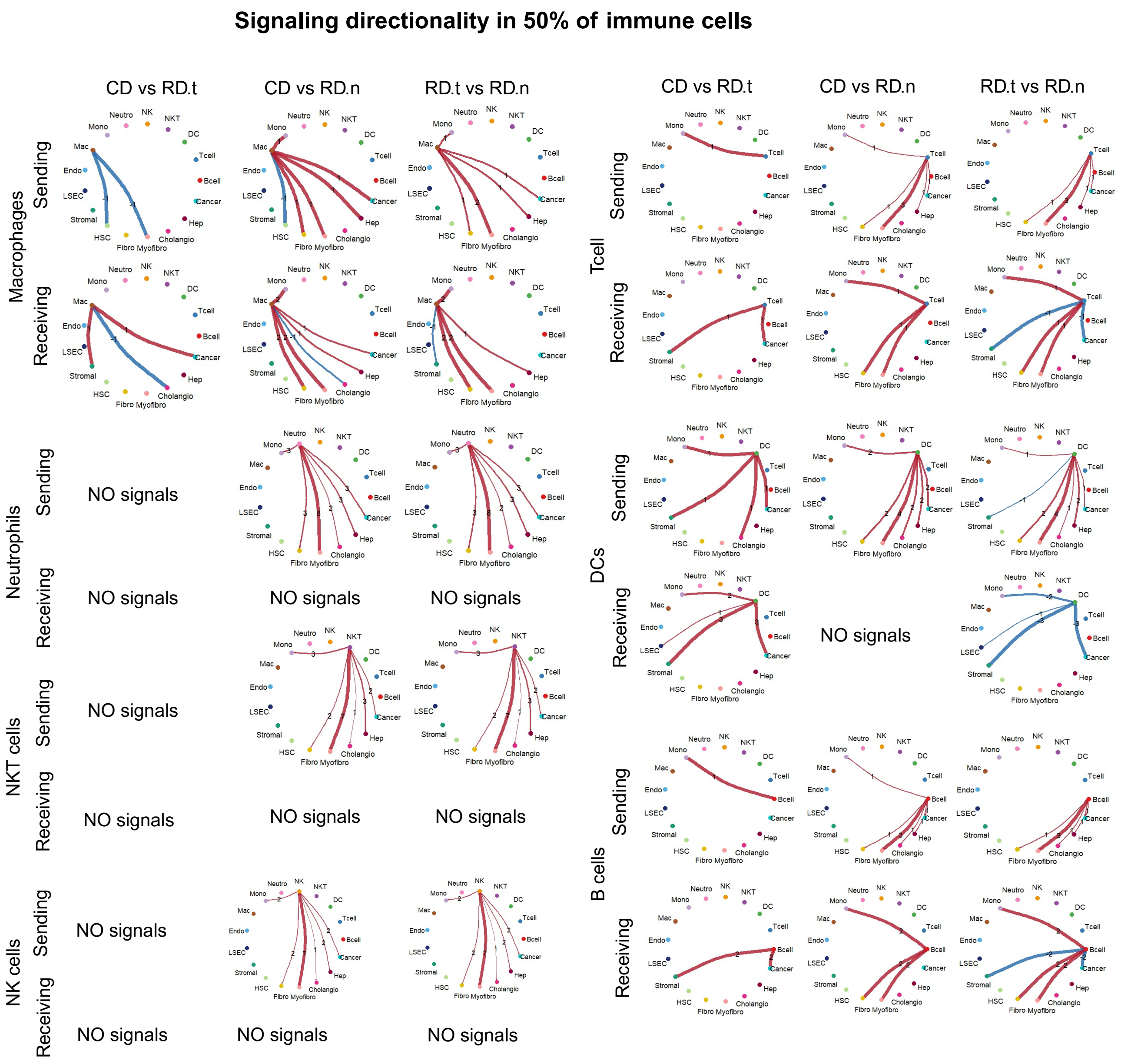


**Figure S12.** The 50% thresold analysis showing the differential number of interactions sent (upper) and received (lower) by Macrophages, Neutrophils, NKT cells, NK cells, T cells, DCs, and B cells across comparisons CDvRD.t (left columns), CDvRD.n (middle columns), RD.tvRD.n (right columns). Red arrows depict increased number of signaling events and blue arrows show decreased events in the RD.t group compared to CD (CDvRD.t), RD.n group compared to CD (CDvRD.n), and RD.n group compared to RD.t (RD.tvRD.n). No signals means there was no difference in the number of signaling events.

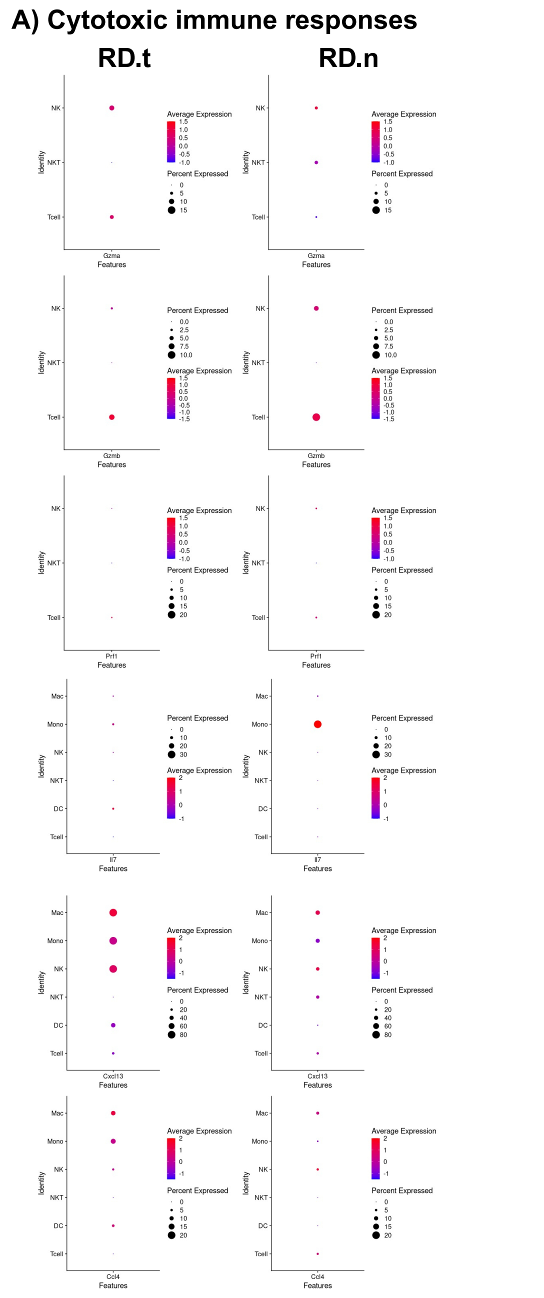


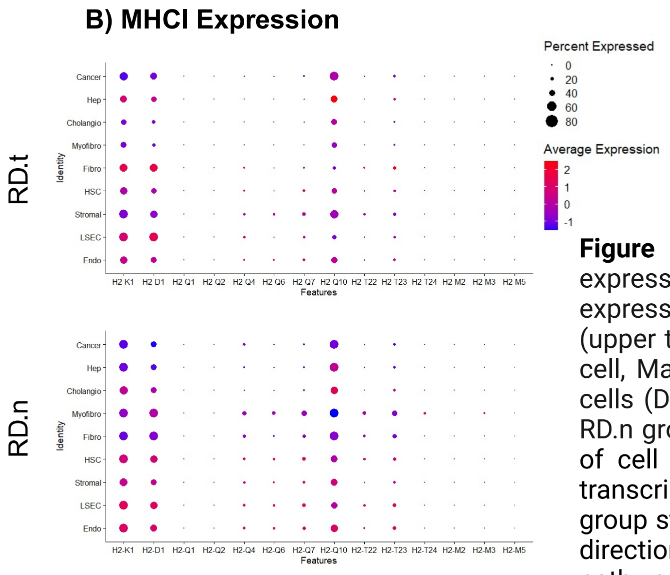


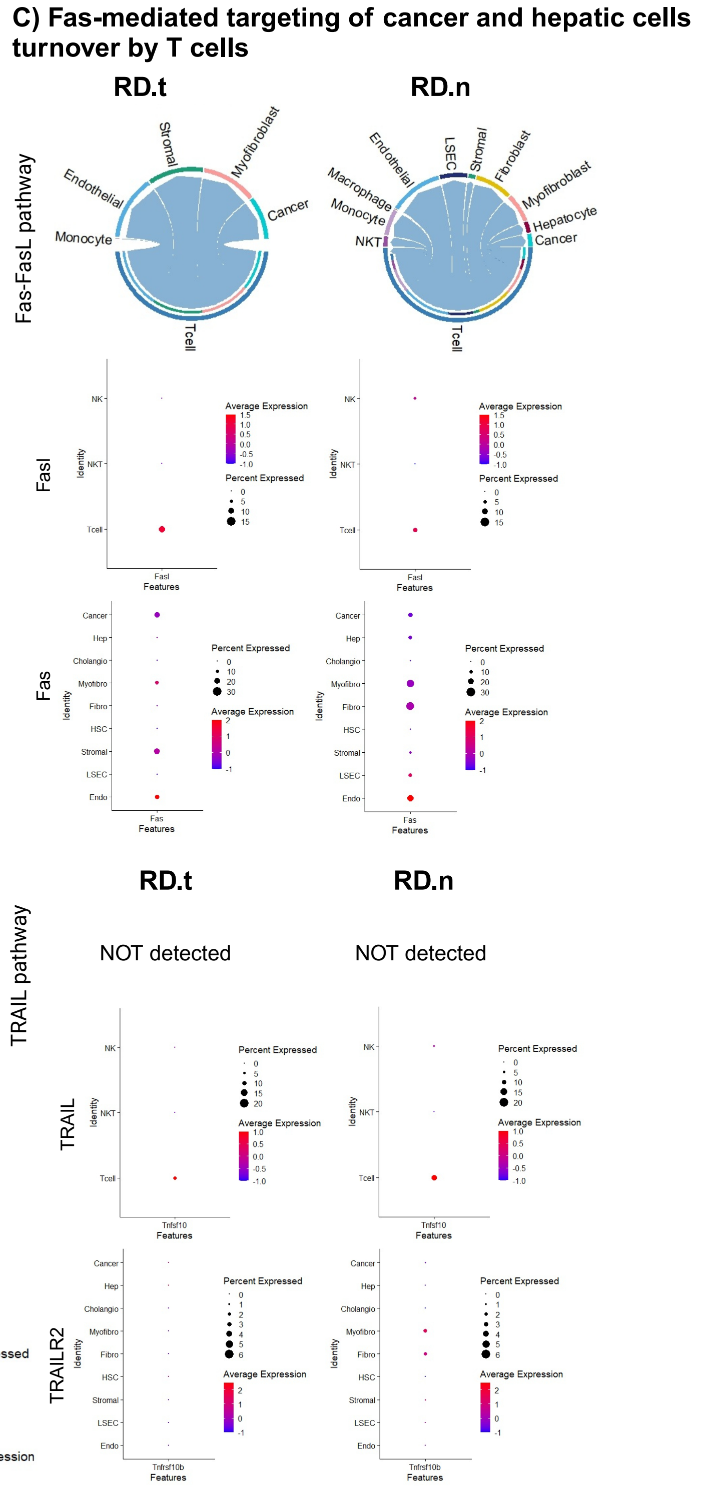


**Figure S13**. A) DotPlots showing percent cell population expression and log-normalized average transcript expression of Gzma, Gzmb, and Prf1 in NK, NKT, and T cells (upper three panels), and Cxcl13, Ccl4, and IL7 in NK, NKT, T cell, Macrophages (Mac), Monocytes (Mono), and Dendritic cells (DC) (lower three panels) in the RD.t (left column) and RD.n groups (right column). B) DotPlots showing percentage of cell population expression and log-normalized average transcript expression of MHCI molecules in RD.t and RD.n group structural cells. C) Chord diagrams depicting signaling directionality detected in 5% threshold analyses for cytotoxic pathways (TRAIL and Fas) in the RD.t and RD.n groups. DotPlots showing percentage of cell population expression and log-normalized transcript expression of FasL (Fasl, top row), Fas (second row), TRAIL (Tnfsf10, third row), TRAILR2 (Tnfrsf10b, bottom row), in the RD.t and RD.n groups
